# Quantitative single-cell live imaging links HES5 dynamics with cell-state and fate in murine neurogenesis

**DOI:** 10.1101/373407

**Authors:** Cerys S Manning, Veronica Biga, James Boyd, Jochen Kursawe, Bodvar Ymisson, David G Spiller, Christopher M Sanderson, Tobias Galla, Magnus Rattray, Nancy Papalopulu

## Abstract

During embryogenesis cells make fate decisions within complex tissue environments. The levels and dynamics of transcription factor expression regulate these decisions. Here we use single cell live imaging of an endogenous HES5 reporter and absolute protein quantification to gain a dynamic view of neurogenesis in the embryonic mammalian spinal cord. We report that dividing neural progenitors show both aperiodic and periodic HES5 protein fluctuations. Mathematical modelling suggests that in progenitor cells the HES5 oscillator operates close to its bifurcation boundary where stochastic conversions between dynamics are possible. HES5 expression becomes more frequently periodic as cells transition to differentiation which, coupled with an overall decline in HES5 expression, creates a transient period of oscillations with higher fold expression change. This increases the decoding capacity of HES5 oscillations and correlates with interneuron versus motor neuron cell fate. Thus, HES5 undergoes complex changes in gene expression dynamics as cells differentiate.

## Introduction

During embryogenesis cells balance proliferation with differentiation to make cell state transitions that lead to the formation of functional organs. This is exemplified by development of the central nervous system, which requires the balance of neural progenitor maintenance with differentiation during multiple waves of differentiation in to neuronal and glial cell-types^1^. In the dorso-ventral (D-V) axis of the spinal cord elegant experiments have shown that fate decisions require integration of a wide range of signals over time, many in the form of morphogen gradients, resulting in downstream gene expression changes^2,3^.

Single-cell transcriptomics have greatly enhanced our understanding of gene expression changes and networks involved in fate decisions and of the bifurcation points where decisions are made^4–7^. However advances in single-cell live imaging of gene expression have shown that it is often highly dynamic, suggesting that the control of cell state transitions is more complex^8–10^. Rather than being in an on or off state, a handful of transcription factors have been shown to oscillate with periodicity of a few hours^9,11^. Oscillations have been long described in somitogenesis^12^, but are a relatively recent discovery in neurogenesis. This is because unlike somitogenesis where oscillations are synchronous within each somite, they tend to be asynchronous in neural progenitor cells (NPCs) and so required unstable reporters and single cell imaging to be discovered^13^. Thus, it is not only changes in gene expression levels that are important, but the short term dynamics of gene expression can also carry important information for cell state transitions. Indeed, there is experimental and theoretical evidence that cell fate transitions may be controlled by a change in the dynamic pattern of gene expression, which could be from oscillatory to stable expression, or to oscillatory with different characteristics^9,14,15^.

In the case of the transcriptional repressor HES1, a key target of Notch signaling, it has been known that oscillatory expression is driven by transcriptional autorepression coupled with delays, instability of mRNA and protein and non-linearity of reactions, common principles of many biological oscillators^16,17^. Like HES1, HES5 is a Notch target bHLH transcription factor (TF) which is highly expressed by NPCs and decreases in expression as differentiation proceeds^18,19^. Knock-out mice and overexpression studies have shown that HES5 functions to maintain the undifferentiated progenitor state through repression of proneural genes, such as *Neurog2 and Atoh1* that promote neuronal differentiation^20–22^. Like HES1, HES5 has been reported to oscillate in NPCs in vitro^9^.

Changes in HES1 dynamics are mediated by a change of the parameters or initial conditions of the oscillator, likely through changes in mRNA stability or protein translation under the influence of a microRNA, miR-9^23–25^. Other theoretical studies provide additional support for the importance of a change in dynamics by showing that gene expression networks in the D-V dimension of the spinal cord can generate multi-way switches (stable or oscillatory)^26^.

An additional revelation of single-cell live imaging studies is that gene expression is characterised by varying degrees of noise due to the stochastic nature of transcription^27–29^. Current ideas for the role of such embedded stochasticity include cases where it would be an advantage^30,31^ or conversely, an impediment for cell fate decisions^32,33^ and mechanisms to suppress noise after a fate-decision^34^.

However, although these studies have shed new light into the problem of cell-state transition, how cells make decision in the context of multicellular tissue is poorly understood. This is because both single-cell transcriptomics and live imaging data are routinely performed in single cells taken out of the tissue environment. Existing studies of oscillatory expression in the mouse brain and spinal cord lack the statistical power needed to give a comprehensive understanding of the dynamics in the tissue^11,35^. A study using electroporation of a promoter reporter of *Hes5-1* in chicken spinal cord tissue reported activation of Notch signaling throughout the progenitor cell cycle but most frequently before mitosis^36^. However, this approach suffered from plasmid loss and varying degrees of plasmid transfection and did not report on endogenous HES5.

Here, we have developed ex-vivo slice culture of embryonic Venus::HES5 knock-in mouse spinal cord (E10.5) to study the expression dynamics of HES5 in the context of a tissue, with single cell resolution. We report that HES5 expression has a 10-fold range between cells in a single expression domain that arises from short-term fluctuations and longer-term trends of decreasing HES5. We use hierarchical clustering to define distinct clusters of single cell HES5 expression dynamics. New statistical tools show that oscillatory HES5 is more frequently observed in cells that transition towards differentiation where it is coupled with an overall decrease in HES5 expression generating larger instantaneous fold changes. Oscillatory decline of HES5 correlates with interneuron fate, suggesting the dynamics are decoded in the choice of cell fate. By contrast, dividing NPCs are less frequently periodic but significantly more noisy in their HES5 expression. Computational modelling with stochastic differential delay equations, parameterised using experimental values and Bayesian inference, suggest that in the spinal cord tissue environment the *Hes5* genetic oscillator operates close to a bifurcation point where noise can tip it from aperiodic to periodic expression. Taken together, our findings suggest that single progenitor cells in a tissue are noisy and are thus primed to enter a transient oscillatory phase as the cells differentiate. Additionally, our study shows that tissue level single-cell heterogeneity has a complex origin in both short and long term dynamics and that the dynamics are decoded en route to differentiation where they correlate with the choice of cell fate that the cells adopt.

## Results

### Venus::HES5 reporter recapitulates endogenous features

We characterised the Venus::HES5 knock-in mouse^9^ to ensure that it is a faithful reporter of the un-tagged gene. In transverse sections of E10.5 spinal cord Venus::HES5 shows a broad ventral and a smaller dorsal domain (Fig. 1a). The ventral domain, which is the focus of this study, encompasses mainly ventral interneuron (p0-p2) and ventral motor neuron progenitors (pMN) (Supplementary Fig. 1a,b). HES5 is expressed in NPCs and declines in neuronal cells (Fig. 1b), consistent with reports of endogenous HES5^7^.

**Figure 1.**
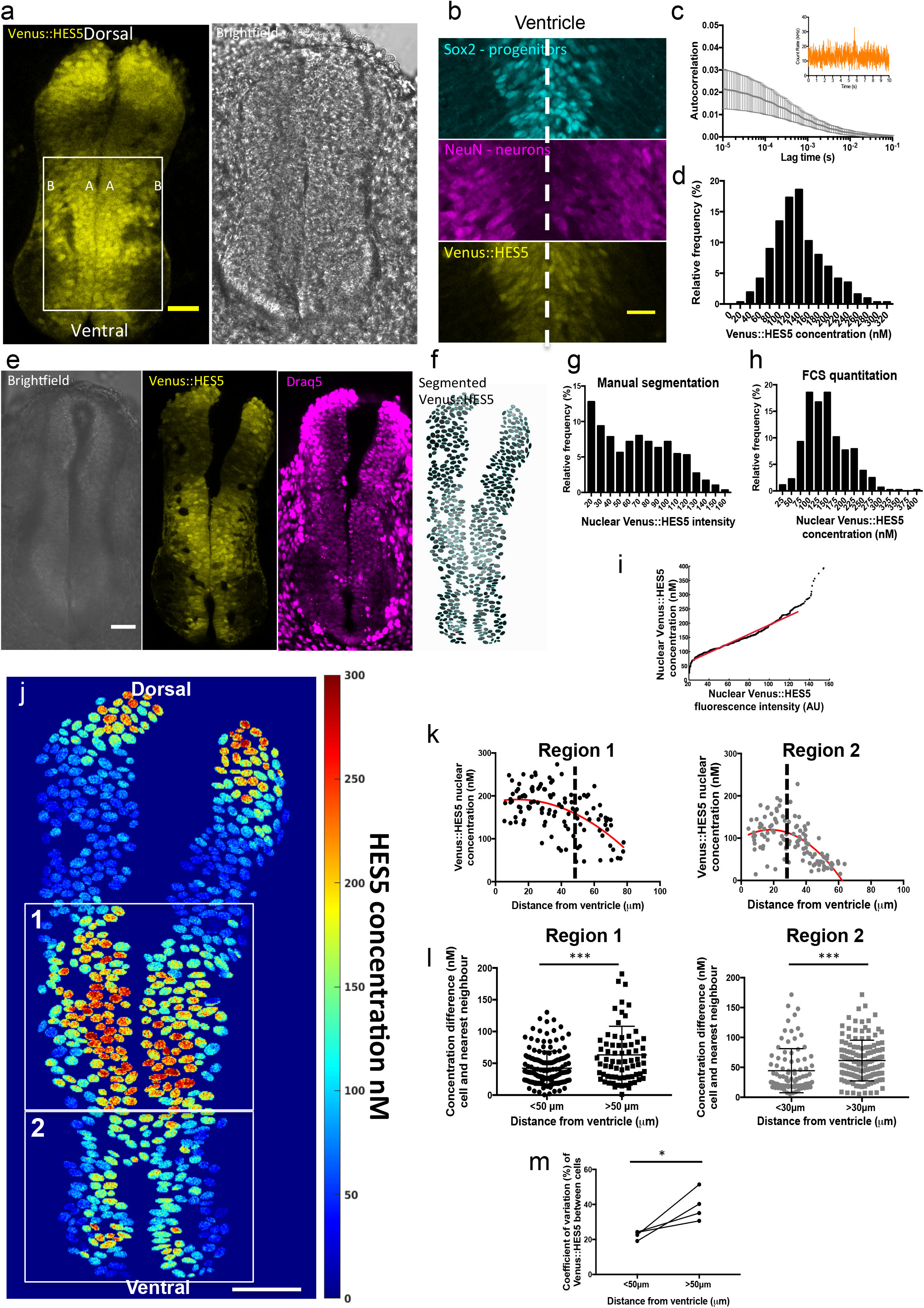
A quantitative transverse map of HES5 expression in the spinal cord. **a**) Transverse slice of live Venus::HES5^+/+^ knock-in mouse spinal cord E10.5 ex vivo. Box identifies ventral domain, A-apical, B-basal. Scale bar 50μm. **b**) Immunofluorescence of E10.5 Venus::HES5 transverse slice of spinal cord. SOX2 - progenitors, NeuN - neurons and endogenous Venus::HES5 signal. Scale bar 30μm. **c**) Average FCS autocorrelation curve. 315 cells in ex-vivo E10.5 Venus::HES5^+/+^ spinal cord ventral region. Inset - example fluorescence count rate from single point within a nucleus. **d**) Nuclear Venus::HES5 concentration in E10.5 Venus::HES5^+/+^ ventral domain. 315 cells, 4 experiments. Mean=140nM, SD=52nM. e) Transverse slice of live Venus::HES5^+/+^ mouse spinal cord E10.5 ex vivo. Draq5 nuclear stain. Scale bar 200μm. **f**) Regions of interest from nuclear segmentation of **e**) with grayscale Venus::HES5 intensity. **g**) Nuclear Venus::HES5 intensity (a.u) in a single live ex-vivo E10.5 Venus::HES5^+/+^ transverse slice (e). n=586 cells. Mean=61a.u SD=39a.u **h**) Nuclear Venus::HES5 concentration in E10.5 Venus::HES5^+/+^ embryos across entire spinal cord. n=442 cells, 4 experiments. Mean=148nM, SD=58nM. **i**) Quantile-quantile plot of nuclear Venus::HES5 concentration (h) vs nuclear Venus::HES5 intensity (g) for E10.5 homozygous embryos. Red line - linear fit over middle 90% range. **j**) Quantitative map of nuclear Venus::HES5 concentration in whole live E10.5 spinal cord. Colour bar shows Venus::HES5 concentration by scaling intensity values according to linear fit of Q-Q plot in **i**). Scale bar 50μm. **k**) Nuclear Venus::HES5 concentration by distance from ventricle in region 1 (upper box in **j**) and region 2 (lower box in **j**) **l**) Concentration difference between a cell and its nearest neighbours for cells less than or greater than 50μm (region 1) from the ventricle (n=154, n=73 cells respectively. p=0.0007 (***) in Mann-Whitney test), or 30μm (region 2) from the ventricle (n=91, n=135 cells respectively. p<0.0001 (***) in Mann-Whitney test). **m**) Coefficient of variation in Venus::HES5 intensity between cells less than or greater than 50μm from the ventricle in ventral domain in E10.5 Venus::HES5 embryos. (n=4 embryos, at least 24 cells per embryo, 2 experiments, p=0.04 (*) in paired t-test.). Error bars – SD. Source data are provided in a Source Data file.

Both mRNA and protein half-lives of Venus::HES5 are unstable with similar values to untagged HES5 (approximately 30 mins for the mRNA and 80-90 mins for the protein). These findings confirm that the Venus::HES5 fusion protein is a faithful reporter of endogenous un-tagged HES5 expression (Supplementary Fig. 1c-f).

### Quantification of range and level of HES5 expression

Dynamic expression can give rise to tissue level single-cell heterogeneity which may be masked by population averaging. Here we use absolute quantitation of Venus::HES5 molecules at the single cell level by Fluorescence Correlation Spectroscopy (FCS) in live homozygous Venus::HES5 E10.5 embryo slices (Fig. 1c,d Supplementary Fig. 2a-d). FCS is an absolute quantification method that records fluorescence emitted as molecules diffuse through a minute volume^37^. The temporal correlation of the signal over time is indicative of the number of molecules present and their diffusion characteristics. Ssing FCS on wild-type E10.5 spinal cord tissue we confirmed that unlike intensity-based techniques FCS count-rate was minimally affected by auto-fluorescence, (Supplementary Fig. 2b). In Venus::HES5^+/+^ embryos single cells showed a 10-fold range of nuclear Venus::HES5 protein expression within the ventral Venus::HES5 expression domain, from 26nM to 319nM. (Fig. 1d). The mean Venus::HES5 nuclear concentration was calculated as 140nM, or 46,250 molecules per nucleus. Heterozygous embryos showed lower mean protein expression, as could be expected by monitoring the expression of one allele (Supplementary Fig. 2e). These findings show a high degree of variability in Venus::HES5 expression between cells which is similar in homozygous and heterozygous embryos suggesting that integrating the expression from 2 alleles does not diminish the variability that cells experience.

### Quantitative map of HES5 expression heterogeneity

FCS can be performed for a limited number of live cells in the tissue, while an intensity map of the Venus signal can be obtained for all cells from snapshot images. We combined the two approaches^38^ by plotting the distribution of single-cell Venus::HES5 intensities from manual segmentation of nuclei in a single slice (Fig. 1g) against the distribution of single-cell FCS protein concentration (Fig. 1h) over multiple slices and experiments. The resulting quantile-quantile (Q-Q) plot was linear and only deviated from linearity at the very high and low values (Fig. 1i). We therefore translate intensity in an image into protein concentration (Fig. 1j) by scaling the intensity value by the gradient of the linear Q-Q plot. Once the Venus::HES5 concentration distribution has been obtained it can be applied to multiple images to generate more quantitative maps without needing to repeat the FCS (Supplementary Fig. 2f,g).

We used the quantitative map to investigate global and local patterns of HES5 concentration. We split the ventral domain into 2 regions due to the difference in width of the ventricular zone along the D-V axis (indicated by boxes in Fig. 1j) We observed a non-linear global reduction of Venus::HES5 concentration with increasing distance from the ventricle (Fig. 1k). The shoulder-point corresponded to around 50μm and 30μm in the dorsal-most (1) and ventral-most (2) regions respectively, suggesting that at this distance, cells start to decrease HES5. At any given distance there is large cell-to-cell variability in Venus::HES5 concentration. The concentration difference between a cell and its nearest neighbour (Supplementary Fig. 2h) increased further away from the ventricle, reaching a maximum of 191nM, a 4.5-fold difference (Fig. 1l). This trend was confirmed in embryos that had not undergone intensity:concentration scaling (Fig. 1m). Thus, further from the ventricle a global reduction of Venus::HES5 expression is accompanied by increasing heterogeneity.

### Clustering indicates distinct Venus::HES5 expression dynamics

Single cell expression heterogeneity may be the result of multiple possibilities: i) fluctuating expression alone (Fig. 2b), which could be periodic and asynchronous ii) distinct but stable cell-state subpopulations (Fig. 2c) or iii) an expression decline as cells transition from one stable state to another (Fig. 2d). Hypothesis (i) implies HES5 expression satisfies ergodicity, i.e. variability in a single cell over time can recapitulate the tissue level heterogeneity^39^. To resolve potential mechanisms that generate heterogeneity, we performed live imaging of Venus::HES5 expression dynamics in ex-vivo slices. We used tamoxifen-dependent recombination in SOX1 + cells 18hrs prior to imaging to label NPCs or cells of neuronal progeny with H2B::mCherry (Fig. 2e,f).

**Figure 2.**
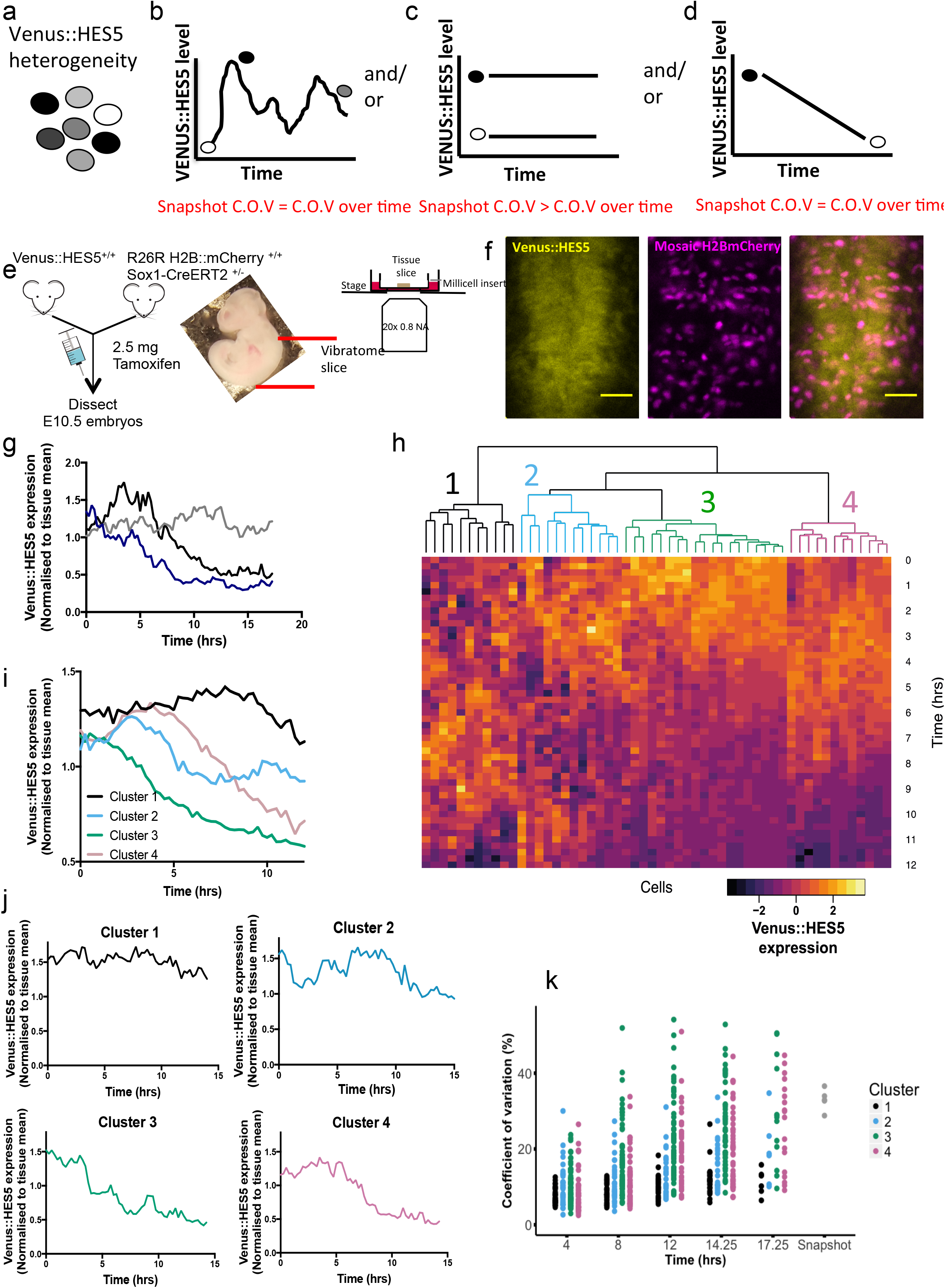
Clustering indicates distinct Venus::HES5 expression dynamics. **a**) Schematic of snapshot Venus::HES5 intensity variability and **b-d**) possible non-mutually exclusive causes. **b**) Stable sub-populations of cells have different expression levels. **c**) single state of cell can traverse all intensity levels **d**) cells undergo one-way transition from high to low levels of expression. **e)** Schematic of experimental approach to image Venus::HES5 expression dynamics from a single endogenous locus. **f**) Snapshot of ex-vivo live E10.5 Venus::HES5 Sox1Cre:ERT2 Rosa26RH2B::mCherry spinal cord slice culture. Scale bar 40μm. **g**) Example single cell traces of normalised Venus::HES5 protein expression in ex-vivo live E10.5 heterozygous Venus::HES5 spinal cord slice cultures. Individual H2B::mCherry+ cells were tracked over time in slice cultures. Single cell Venus::HES5 intensity values were normalised to the tissue mean intensity over time. **h**) Representative dendrogram from hierarchical clustering of standardised single cell Venus::HES5 protein dynamics in E10.5 heterozygous Venus::HES5 spinal cord slice culture in 1 experiment. Columns show standardised individual cell Venus::HES5 expression dynamics in a heatmap aligned to start at t=0, the start of tracking. Rows represent time points after start of individual cell tracking. 54 cells tracked for 12-hour time window with 15-minute frame intervals. **i**) Mean Venus::HES5 expression dynamics for cells in each cluster in a representative experiment corresponding to dendrogram in e). (Cluster 1 - 11 cells, cluster 2 - 11 cells, cluster 3 - 21 cells, cluster 4 - 11 cells). **j**) Example single cell traces for each cluster of normalised Venus::HES5 expression in ex-vivo live E10.5 spinal cord slice cultures. **k**) Left - coefficient of variation (C.O.V) of single-cell Venus::HES5 expression over time within 4, 8, 12,14 and 17.25 hour windows. Cluster 1 – black, cluster 2 – sky blue, cluster 3-green, cluster 4 – pink. 181 cells, 3 experiments clustered separately, single points show C.O.V from a single-cell timeseries. Right - C.O.V in Venus::HES5 protein levels between cells measured at a single time point. 5 ex-vivo E10.5 Venus::HES5 slices in 2 experiments, single points show COV between cells in a single slice. Source data are provided in a Source Data file.

We observed multiple types of single-cell Venus::HES5 dynamic behaviours in heterozygous cells (Fig. 2g) over a time period of 12-15 hours. Hierarchical clustering of the standardised Venus::HES5 intensity timeseries suggested 4 clusters of longterm Venus::HES5 expression dynamics (Fig. 2h and Supplementary Fig. 4a,b). Cells in cluster 1 and 2 showed fluctuating expression around a stable mean whereas cells in clusters 3 and 4 showed gradually decreasing and fluctuating HES5 expression (Fig. 2h). The non-standardised mean expression of cells in each cluster maintained this trend (Fig. 2i) which is further exemplified by single cell traces (Fig. 2j).

The coefficient of variation (C.O.V, standard deviation of intensity divided by the mean intensity) of Venus::HES5 over time in single cells increased over 4,8,12,14.25 and 17.25 hours (Fig. 2k). By 8-12 hours multiple cells in clusters 3 and 4 had reached similar or higher levels of variation as the variation observed between cells at a single snapshot (Fig. 2h) suggesting that declining expression is a major contributor to the tissue heterogeneity. In contrast, cells in clusters 1 and 2 rarely reached tissue-levels of variation between cells, suggesting that short-term dynamics have a lesser contribution to overall tissue heterogeneity and excluding scenario in Fig. 2b. Thus heterogeneity is generated by a mix of declining expression (long-term trends, scenario Fig. 2d,) and dynamic fluctuations (short term dynamics) around a slowly varying mean.

### Venus::HES5 expression dynamics correlate with cell-states

We hypothesise that the different clusters of Venus::HES5 expression may represent different cell-states. It is well known that proliferating NPCs (SOX1+/2+) are found apically in the ventricular zone, undergo inter-kinetic nuclear migration (INM) dividing at the apical surface^1,40^. Newly born cells fated towards neuronal differentiation migrate basally away from the apical surface, exit the cell cycle and turn on markers of differentiation (Tuj1 and NeuN)^40^. We therefore sought to infer cell state by position, motility and division of cells using progenitor/neuronal immunochemistry data in ex-vivo slices as reference.

The average position of cells in cluster 1 was significantly closer to the ventricle than those in cluster 3 (Fig. 3c). Further, in a zone greater than 50μm from the ventricle very few cells of cluster 1 reside and cells in cluster 3 are more abundant (Fig. 3d). By contrast, the zone within the first 50μm of the ventricle is equally occupied by cells in clusters 1-4 (Fig. 3d).

**Figure 3.**
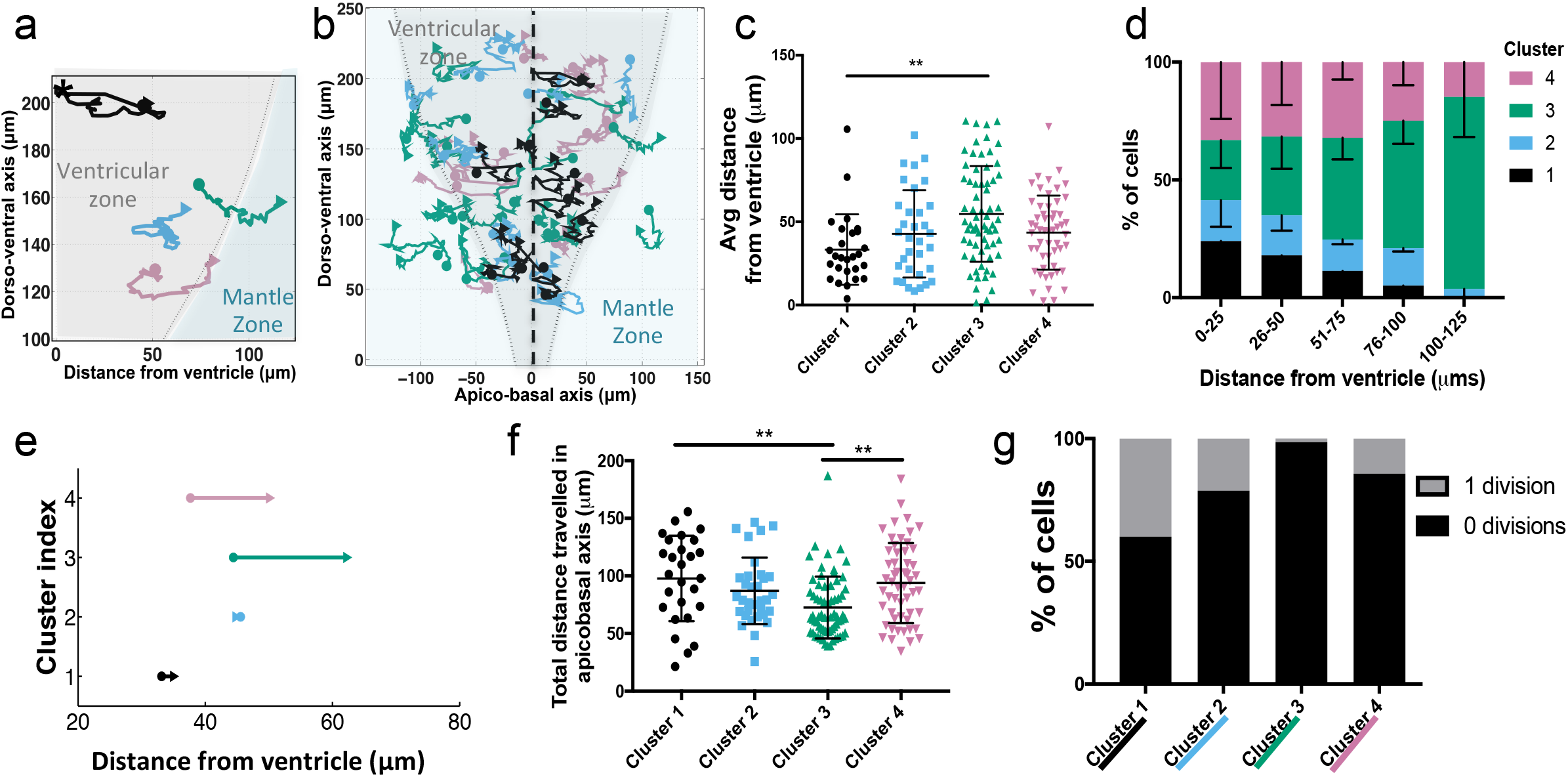
Single cell Venus::HES5 dynamics correlate with cell state. **a**) Example single cell tracks for each cluster from 1 representative experiment. ‘*’ denotes cell division. Dorso-ventral axis in μm from floorplate. **b**) Single cell tracks (n=54) over 12-hours in a single E10.5 spinal cord slice movie. Black dotted line - ventricle. Grey - ventricular zone and green - mantle zone measured by Sox2/NeuN immunostaining of 2 ex-vivo slices. **c**) Average distance of single cell from ventricle over 12-hour track. Cells from 3 experiments clustered separately. Kruskall-Wallis with Dunn’s multiple comparisons test indicated cluster 1 vs 3 adjusted p-value = 0.0014 (**) **d**) Percentage of any cells found in 25μm windows from ventricle in each cluster. 2-way ANOVA with Tukey multiple comparison test shows no difference between clusters <50μm from ventricle. 76-100μm from ventricle cluster 3 vs 1 p<0.0001, 3 vs 2 p=0.0009, 3 vs 4 p=0.014. **e**) Displacement of cells in each cluster. Dot - average start position, arrow - average finish position. **f**) Total distance travelled of single cells. Line is mean with SD. Kruskal-Wallis test with Dunn’s multiple comparison test shows cluster 1 vs 3 adjusted p=0.003 (**), cluster 3 vs 4 adjusted p=0.0017 (**). **g**) Percentage of cells per cluster undergoing 0 or 1 divisions in 12 hours. Chi-squared test of frequency data p=0.0002. 181 cells, 3 experiments clustered separately. Cluster 1 - black (n=27 cells), cluster 2 - sky blue (n=33 cells), cluster 3 - green (n=67 cells), cluster 4 - pink (n=54 cells). Error bars – SD. Source data are provided in a Source Data file.

Nuclei of cells in cluster 1 moved both apically and basally, consistent with inter-kinetic nuclear migration (INM) but had the shortest displacement as they returned apically. Meanwhile nuclei of cells in cluster 3 and 4 had a larger displacement which was unidirectional towards the basal side (Fig. 3a,b,e,f, Supplementary Fig. 5a and Supplementary Movie 1) suggesting they are on their way to differentiation. Immunostaining and measurement of the SOX2+ domain showed that many cells in cluster 3 and 4 moved out from the SOX2+ zone into the mantle zone with concurrent decreasing Venus::HES5 (Fig.3a,b, Supplementary Fig. 5b,c,d). H2B::mCherry dynamics did not decrease in level, neither close nor far away from the ventricle, and were similar between clusters (Supplementary Fig. 4c,5e).

Cells divided at the apical surface (Supplementary Fig. 5f) and the number of divisions was significantly higher in cluster 1 and 2; indeed, very few cells in cluster 4 and no cells in cluster 3 were observed to divide (Fig. 3g). Given these findings, we inferred that cells in cluster 1 and 2 are proliferating progenitors and cells in cluster 3 and 4 are transitioning towards differentiation.

What are the differences between cluster 1 and 2 and between clusters 3 and 4? We inferred cell-cycle phase based on cell position and trajectory and we found no difference in the cell cycle profiles between cells in cluster 1 and 2 (Supplementary Fig. 5g). Cells in cluster 1 have higher levels of Venus::HES5 than cells in cluster 2 when levels are normalized for z-depth of the cell into the tissue (Supplementary Fig. 5h). Cells in cluster 4 tend to be delayed in the decrease in Venus::HES5 levels compared to cells in cluster 3 (Fig. 2i) and show a small total number of divisions (14%; Fig. 3g), in contrast to 1.5% of divisions in cluster 3.

We confirmed our interpretation of cell-state by using the Notch inhibitor DBZ to promote differentiation^7^. Spinal cord ex-vivo slices treated with 2μM DBZ showed significantly lower mean Venus::HES5 intensity than control (Fig. 4a, Supplementary Fig. 6a) and an increase in the early neuronal marker β-tubulin especially in apical regions (Fig. 4b). The disorganisation of the neural tube in DBZ treated slices is similar to *Hes* KO phenotypes^41^, consistent with *Hes5* being a downstream target of Notch. The average position of single cells in DBZ treated slices was further from the ventricle (Supplementary Fig. 6b) and they showed significantly increased apico-basal displacement confirming that Notch inhibition had pushed cells towards basal migration and differentiation (Fig. 4c). Hierarchical clustering of standardised Venus::HES5 single-cell intensities showed that 98% of cells in the DBZ treated slices were found in clusters 3 and 4 (Fig. 4d,e). Specifically, the timing of Venus::HES5 decline, the COV of Venus::HES5 over time and the number of divisions is consistent with most of the DBZ treated cells falling into cluster 4-type dynamics (Supplementary Fig. 6c,d,e), while the distribution of control DMSO Venus::HES5 cells recapitulated the presence of all 4 clusters (Supplementary Fig. 6f,g).

**Figure 4.**
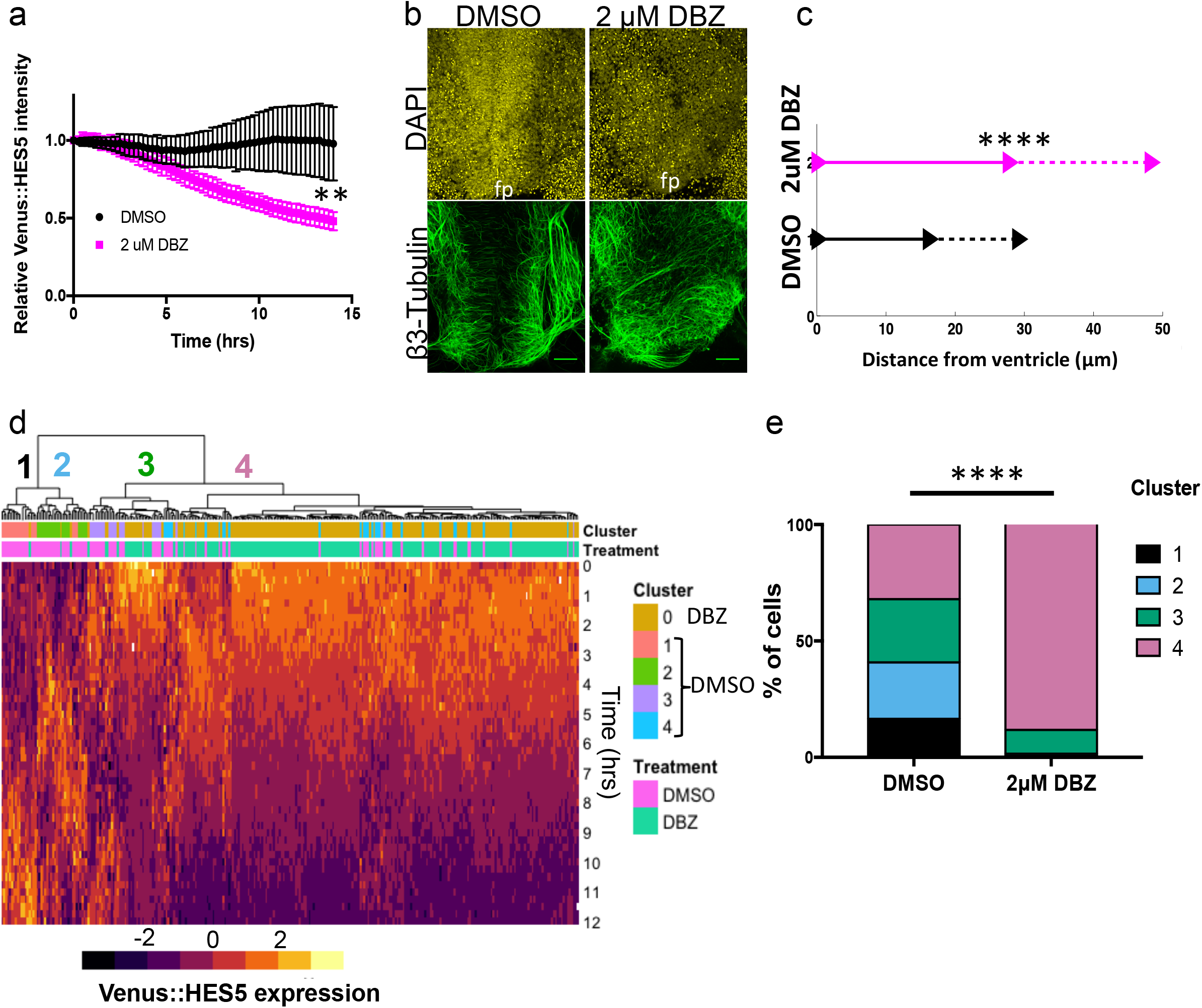
Notch inhibition enriches for cluster 4-type Venus::HES5 dynamics. **a**) Relative Venus::HES5 intensity of E10.5 ex-vivo slices cultured in control DMSO (black) or 2μM DBZ (pink). Error bars - SD (3 experiments). Endpoint intensity - Mann Whitney two-tailed test p=0.0095 (**). **b**) β-III tubulin (early neuronal marker) staining of DMSO or 2μM DBZ treated E10.5 ex-vivo slices. fp – floorplate. Scale bar 70μm. **c**) Displacement of single cells away from ventricle in apico-basal axis in control DMSO or 2μM DBZ treated E10.5 slices. Bold line - average, dashed line - SD from 3 experiments. Two-tailed t-test p<0.0001 (****). **d**) Hierarchical clustering of standardized single-cell Venus::HES5 expression in DMSO and 2μM DBZ treated E10.5 ex-vivo slices. Columns show standardised individual cell Venus::HES5 expression dynamics in a heatmap. Rows represent time points after start of individual cell tracking. Cells tracked for 12-hour time window with 15-minute frame intervals. 295 cells, 3 experiments clustered together. Cluster labels defined using clustering of DMSO alone. Pink – DMSO cells, green - DBZ cells. **e**) Percentage of cells in each cluster in DMSO and 2μM DBZ treated E10.5 ex-vivo slices. Frequency data subject to Chi-squared test showed p<0.0001 (****). DMSO n=100 cells, 2μM DBZ n=195 cells, 3 experiments. Source data are provided in a Source Data file.

We conclude that cells characterised by a temporally fluctuating Venus::HES5 expression pattern around a high mean (cluster 1 and 2) are proliferating NPCs maintained by Notch signalling, while cells with decreasing Venus::HES5 levels over time (clusters 3 and 4) are neural cells undergoing cell state transition to differentiation. We do not know the significance of the subtle differences between clusters 1 and 2 or 3 and 4 but we suggest that the simplest interpretation of our data is that clusters 1+2 give rise to clusters 3+4. However more complex alternatives may exist, such as subtle heterogeneity in progenitors translated linearly to neuronal progeny heterogeneity.

### Differentiating cells are oscillatory and progenitors noisy

Previous reports show periodic HES5 expression in embryonic mouse cortical NPCs using luciferase and fluorescence imaging^9^ but statistical analysis has not been performed. Here, we have focused on the endogenous Venus::HES5 fluorescent fusion protein because unlike luciferase, it allows single cell spatial resolution in the tissue environment by confocal microscopy. The t_1/2_ of Venus maturation (15 mins)^42^ is suitably short compared to HES5 protein half-life (80-90 mins, Supplementary Fig. 1d). HES5 traces show a high degree of variability (Supplementary Fig. 7) and detecting oscillatory gene expression in such noisy timeseries, is challenging. We have previously developed an approach for statistical determination of oscillations in noisy bioluminescent data^43^. Here, we extend this method to take into account that fluorescence intensity timeseries from tissue are inherently more noisy partly because they do not involve the long integration times associated with Luciferase imaging (Methods).

To analyse oscillations, we first subtracted long-term changes in level (trend) caused by HES5 downregulation (Fig. 5a). We then analysed detrended data with an oscillatory covariance model and inferred the period, amplitude and lengthscale (Fig. 5b). Lengthscale accounts for variability in the peaks over time. We compared the oscillatory (alternative) model fit and aperiodic (null) covariance model fit using the log-likelihood ratio (LLR), which is high for oscillators (Supplementary Fig. 8a) and low for non-oscillators (Supplementary Fig. 8b). Finally, we identified oscillatory cells in each experiment using a strict false-discovery rate criteria set at 3% (Supplementary Fig. 8d).

**Figure 5.**
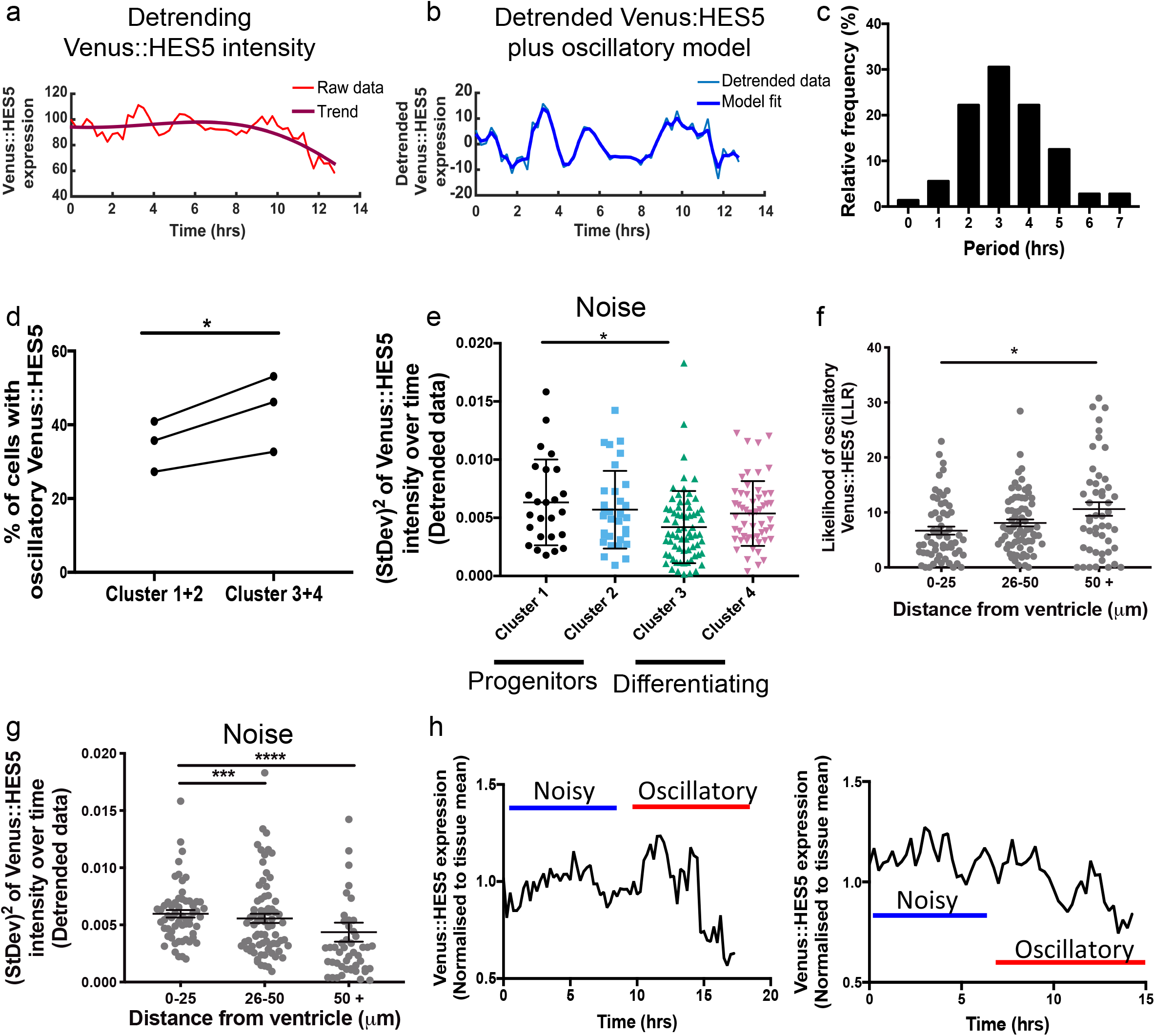
Differentiating cells are oscillatory and progenitors are noisy. Example single-cell traces of Venus::HES5 expression subject to oscillatory test. **a**) Raw single-cell Venus::HES5 intensity timeseries with overlaid long-term trend in bold. **b**) Detrended single-cell Venus::HES5 intensity timeseries with overlaid OUOsc oscillatory model in bold. **c**) Single-cell periods of Venus::HES5 protein dynamics from oscillatory cells. Mean=3.3hours, SD=1.4hours. n=72 cells, 3 experiments. **d**) Percentage of single cells classified as having oscillatory Venus::HES5 protein expression in cluster 1+2 vs cluster 3+4. n = 3 experiments, p-value = 0.04 (*) in Wilcoxon paired test, two-tailed. Single-cell raw Venus intensity was detrended and subject to oscillatory test (see methods). **e**) Noise in single cell Venus::HES5 expression dynamics per cluster as measured by the squared standard deviation of de-trended Venus::HES5 signal over time. Lines show mean, error bars - SD. Kruskal-Wallis with Dunn’s multiple comparison test shows cluster 1 vs 3 adjusted p=0.04 (*). 181 cells, 3 experiments clustered separately. Cluster 1 - black (n=27 cells), cluster 2 - sky blue (n=33 cells), cluster 3 - green (n=67 cells), cluster 4 - pink (n=54 cells). **f**) Likelihood of a cell having oscillatory Venus::HES5 expression indicated by LLR score plotted by average distance of the cell away from the ventricle over 12-hour track. Bars show mean and error bars show SEM. Kruskal-Wallis with Dunn’s multiple comparison test shows 0-25μm vs 50+μm adjusted p=0.03 (*). **g**) Noise in single cell Venus::HES5 expression dynamics plotted by average distance of the cell away from the ventricle over 12-hour track. Bars show mean and error bars show SEM. Kruskal-Wallis with Dunn’s multiple comparison test shows 0-25μm vs 26-50 adjusted p= 0.0007 (***) and 0-25μm vs 50+μm adjusted p=<0.0001 (****). **h**) Example single-cell timeseries of relative Venus::HES5 protein expression in ex-vivo live E10.5 Venus::HES5 spinal cord slice cultures showing noisy to oscillatory transition in Venus::HES5 dynamics. Source data are provided in a Source Data file.

We found that overall 41% of cells in E10.5 spinal cord ex-vivo showed oscillatory Venus::HES5 expression (Supplementary Fig. 9a), while the rest were fluctuating and aperiodic. The mean period of Venus::HES5 oscillations was 3.3±0.3 hours (±S.D) (Fig. 5c) while H2B::mCherry expression from the ROSA26 locus in the same nuclei was aperiodic (Supplementary Fig. 9a).

We also imaged cells dissociated from the spinal cord of heterozygous Venus::HES5 mouse embryos and cultured in vitro, as this matches the experimental set-up used previously^9^ (Supplementary Fig. 9a,b). The occurrence of oscillatory Venus::HES5 expression was higher in dissociated cells compared to cells in the tissue environment (Supplementary Fig. 9a). Nuclear Venus::HES5 concentrations were also significantly lower in dissociated cells (Supplementary Fig. 9d). This finding confirms the ability of our methods to detect oscillations and further suggests that HES5 dynamics are influenced by the tissue environment, although many factors change between in vitro and ex vivo conditions. Interestingly, there was no difference in the percentage of oscillatory cells isolated from heterozygous versus homozygous mice confirming that cells experience oscillatory HES5 dynamics (Supplementary Fig. 9a,d).

We next sought to determine which of the clusters contain cells with oscillatory expression. Oscillations were not restricted to proliferating progenitor cells, instead Venus::HES5 oscillations were more frequently observed in cells on their way to differentiation (clusters 3 and 4) than dividing progenitors in clusters 1 and 2 (Fig. 5d). By contrast, proliferating progenitors in cluster 1 had significantly greater noise than differentiating cells in cluster 3, (noise measured by squared-standard deviation of de-trended Venus::HES5 signal (Fig. 5e)). In agreement with this, the likelihood of a cell to have oscillatory Venus::HES5 significantly increased with an increasing average distance from the ventricle (Fig. 5f), whereas noise decreased (Fig. 5g)

Given that progenitor cells close to the ventricle (cluster 1 & 2) are likely to turn into the transitory and differentiating cells in cluster 3 and 4, we conclude that progenitor cells have high, dynamic and noisy Venus::HES5 expression which evolves in to a more oscillatory signal as Venus::HES5 decreases and the cells undergo differentiation. Although our observation time window is relatively short, data collected from a few cells in cluster 1 demonstrate this noisy to oscillatory transition in Venus::HES5 expression, supporting this view (Fig. 5h, Supplementary Movie 2).

### *Hes5* network poised at aperiodic to oscillatory transition

To understand how the HES5 dynamics of clusters 1 and 2 are generated and how they may transition from aperiodic to periodic expression, we used a stochastic delay differential equation model of an auto-negative feedback network (Fig.6a and Methods)^30,44–46^. This model applies to progenitors in clusters 1 and 2 where HES5 fluctuates around a more or less stable mean. We parameterized the model using protein and mRNA half-lives (Supplementary Fig. 1c,d) and Approximate Bayesian Computation (ABC)^47^ to search for parameters that give rise to experimentally observed summary statistics of HES5 expression (Methods). ABC has advantages over commonly-used point estimates because it provides a probability distribution for estimated parameters thus quantifying parameter uncertainty. We found that the experimentally measured distribution of oscillation periods and relative standard deviation values in clusters 1 and 2 (Supplementary Fig. 10a,b) are consistent with the predictions from this model (Fig. 6b,c).

**Figure 6.**
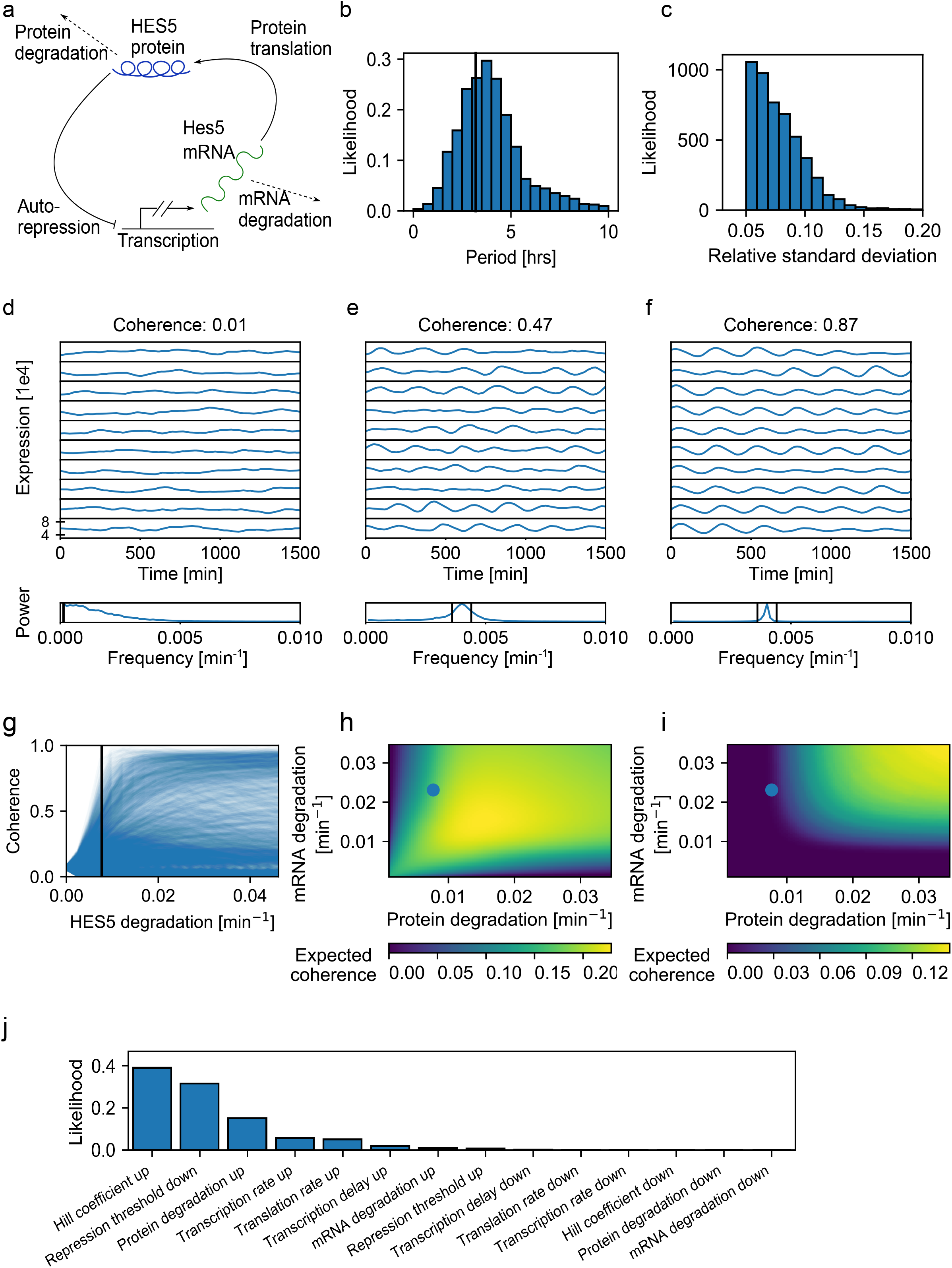
*Hes5* network is poised at aperiodic to oscillatory transition point. **a**) Schematic of stochastic model for genetic autorepression of HES5. **b**) Bayesian posterior model predictions of HES5 periods. Periods are extracted from simulated data of 12h duration using Hilbert transforms. Black line indicates mean of experimentally measured periods. Mean=4.47 hours, SD=2.51 hours, n=4901 samples. **c**) Distribution of model predicted relative standard deviations (standard deviation/mean) of HES5 expression over time. Mean =0.078, SD=0.023, n=4901 samples. The distribution approaches zero around 0.15, the experimentally determined maximum value of standard deviation of Venus::HES5 over time (detrended data) in proliferating progenitors in clusters 1 and 2 (Supplementary Fig. 10b). **d-f**) Ten example traces generated using the model are shown at three different parameter points. The power spectrum does not have a dominant non-zero peak in d) whereas the power spectra in e) and f) do have a dominant non-zero peak with decreasing width from e) to f) showing increasing coherence. Parameter values are (d) α_m_=0.64min^−1^, α_p_=17.32min^−1^, P_0_=88,288.6, T=34min, n=5.59 (e) α_m_=39.93min^−1^, α_p_=21.56 min^−1^, P_0_=24,201.01, T=33min, n=4.78 (f) α_m_=44.9min^−1^, α_p_=3.13min^−1^, P_0_=35,080.2, T=40min, n=5.62. The half-lives of the protein and mRNA are set to 90 and 30 minutes, respectively. **g**) Response curves in coherence when changing the protein degradation rate (n=4901 samples). The black line is located at the degradation rate corresponding to a 90 minutes HES5 protein half-life. **h-i**) Heatmaps showing expected coherence for the stochastic model (**h**) and the deterministic model (**i**) of HES5 expression as protein and mRNA degradation rates are changed. The blue dots mark experimentally measured values for the protein and mRNA degradation rates, corresponding to a 90 and 30 minute half-life, respectively. Experimentally measured degradation rates are located on the slope of increasing coherence values in the stochastic model, and in a region of no expected oscillations in the deterministic model. **j**) Likelihood of inducing oscillations with less than 5hr period from aperiodic fluctuations when changing individual parameters by 50% (n=48503 samples).

HES5 expression simulated from inferred parameters can be aperiodic (Fig. 6d) or oscillatory (Fig. 6e,f) depending on the parameters, as illustrated qualitatively by a sharpening of the peak in the power spectrum and expressed quantitatively by coherence^30^. At unique combinations of parameter values the stochastic model predicts that different proportions of aperiodic and oscillatory HES5 expression will be generated across traces and within the same trace. This is consistent with our experimental observations where less than half of cells pass oscillatory tests and we can observe changes in expression dynamics.

We investigated how HES5 expression may transition from aperiodic to oscillatory in a number of ways. Firstly, we investigated how oscillation coherence varies in response to changing the protein degradation rate across parameter space using Bayesian inference (Fig. 6g where each curve corresponds to one possible parameter combination). The experimentally measured protein degradation rate (protein half-life of 90 minutes, blue-line Fig. 6g) defines a transition point where the range of possible coherence values changes sharply.

We next determined the predicted coherence in relation to the protein and mRNA degradation rates for the full stochastic model (Fig. 6h) and the deterministic model (Fig. 6i). The experimentally measured mRNA and protein degradation rates were located in a region of parameter space where oscillations are expected in the stochastic model, but not in the deterministic model. This is consistent with a full Bayesian comparison between the two (Methods) where the stochastic model is 160 times more likely to describe the HES5 expression statistics (Supplementary Fig. 10d). Our experimentally measured degradation rates predict that the stochastic system is at the boundary of high and low coherence.

Finally we explored which parameters are most likely to generate a change in dynamics between aperiodic and oscillatory HES5. Starting from parameter combinations for which the model predicts aperiodic dynamics, we changed individual model parameters by 50% and recorded the likelihood of this parameter change to induce oscillations (Fig. 6j). This indicated that a range of parameter changes have the potential to induce oscillations, among which increases in the Hill coefficient, decreases in the repression threshold and increases in protein degradation rate are the most likely options.

Taken together, our modelling suggests that the HES5 oscillator in spinal cord NPCs is enabled by noise^30,48^ and operates very close to the boundary between aperiodic and oscillatory model dynamics, where small parameter changes can cause a transition between non-oscillatory (low coherence) and oscillatory (high coherence) expression. It also predicts that increases in the Hill coefficient, and decreases in repression threshold and protein degradation are most likely to initiate oscillatory expression dynamics.

### HES5 oscillations on a downward trend increase fold-changes

Given the higher incidence of oscillatory cells in differentiating cells (cluster 3 and 4) we investigated whether HES5 oscillations are caused by the reduction of HES5 levels. The mean levels between cells is different in clusters 1 and 2, but there was no correlation with the presence of oscillations, arguing against the protein expression level alone having a causative effect for oscillations (Supplementary Fig 11a,b). As expected^49,50^ we did find a positive relationship between Venus::HES5 levels and noise (represented by absolute variance, Supplementary Fig. 11c), which was also captured by the modelling (Supplementary Fig. 11d). Further, treatment with the Notch inhibitor DBZ significantly decreases Venus::HES5 levels, enriches for cluster 4-type dynamics (Fig. 4d,e) but does not significantly change the percentage of oscillators compared to control DMSO within clusters 3 and 4 (Supplementary Fig. 11e and example single cells in Supplementary Fig. 12).

Why then do periodic oscillations occur predominantly during the decay in fluorescence in groups 3 and 4? The maximal peak-to-trough fold change in Venus::HES5 expression, a measure that includes the downward trend, was significantly higher in differentiating cells in clusters 3 and 4 than proliferating progenitors in cluster 1 (Fig. 7a, Supplementary Fig. 11f). Furthermore, within cluster 3, oscillatory cells have a higher mean peak-to-trough fold change than non-oscillatory cells (Fig. 7b,c,d), although differentiating cells eventually undergo amplitude death (Fig. 7e,f). When the declining trend was removed from the data differentiating cells no longer had the increased peak-to-trough changes (Supplementary Fig. 11g). Taken together, these findings suggest that oscillations are combined with a long-term decreasing signal to transiently promote larger fold-changes in HES5 protein than either one alone, forming the decoding phase of the oscillator.

**Figure 7.**
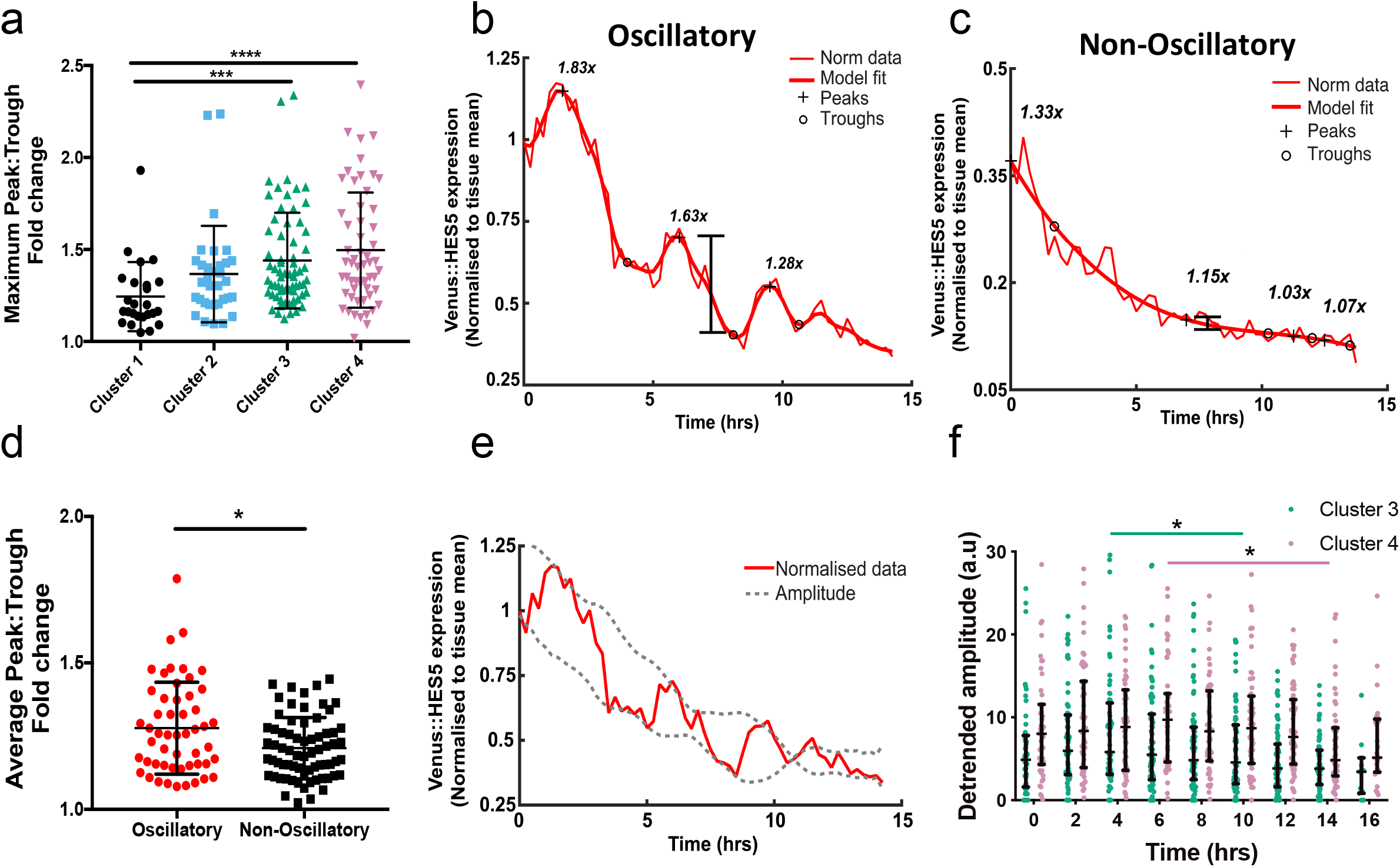
HES5 oscillations on a downward trend increase fold-changes. **a**) Maximum peak-to-trough fold-change in single cell Venus::HES5 expression per cluster. Kruskal-Wallis with Dunn’s multiple comparison test shows cluster 1 vs 3 adjusted p=0.0008 (***), cluster 1 vs 4 adjusted p<0.0001 (****). 181 cells, 3 experiments clustered separately. Cluster 1 (n=27 cells), cluster 2 (n=33 cells), cluster 3 (n=67 cells), cluster 4 (n=54 cells). Peak:Trough fold change is calculated from normalised Venus::HES5 expression including trend. Examples of single-cell Venus::HES5 timeseries in cluster 3 with **b**) oscillatory and **c**) non-oscillatory expression. Bold lines indicate model fit over normalized Venus::HES5 intensity. Plus sign indicates peak and circle indicates trough in intensity values, fold-changes between peak-trough are indicated at relevant peak. **d**) Mean peak-to-trough fold-change in oscillatory (n= 52 cells, 3 experiments) or non-oscillatory (n = 69, 3 experiments) single-cell Venus::HES5 expression in differentiating cells in cluster 3 and 4. p=0.027 (*) in Mann-Whitney test after 2 outliers removed. **e**) Example singlecell timeseries of mean normalized Venus::HES5 expression (red) from cluster 3 showing amplitude death (amplitude indicated by dashed line). **f**) Instantaneous amplitudes from Hilbert transformation of de-trended single cell Venus::HES5 expression observed over time. 121 cells from cluster 3 and 4 in 3 experiments clustered separately. Student’s t-test were used to compare maximum amplitude data in: cluster 3 against subsequent timepoints showing significant decay after 10h p= 0.0470 (*), 12h-16h p<0.0001; cluster 4 showing significant decay after 14h p= 0.0153 (*) and p=0.0195 for 16h. Error bars - SD. Source data are provided in a Source Data file.

### Oscillatory HES5 correlates with interneuron fate

To gain insight into the possible functional significance of oscillatory expression in differentiating cells, we asked whether there is a correlation between oscillatory and non-oscillatory differentiating cells in clusters 3 and 4 and the fate that the cells adopt. Spatial patterning of the ventral spinal cord driven by Shh gradient results in clearly delineated progenitor domains that each give rise to different neuronal subtypes. Therefore distance from the floorplate specifically instructs neuronal sub-type with motor neuron progenitors located more ventrally than interneuron ones. Combining the distance of cells from the floorplate and staining of the cultured ex-vivo slices for motor neuron and interneuron progenitor markers, we found that there is a higher incidence of oscillatory Venus::HES5 expression in differentiating cells that give rise to interneurons than in those giving rise to motor neurons (Fig. 8a-c). We therefore conclude that there are two paths by which HES5 declines, one of which is oscillatory and one which is not, and this correlates well with the fate that these cells adopt.

**Figure 8.**
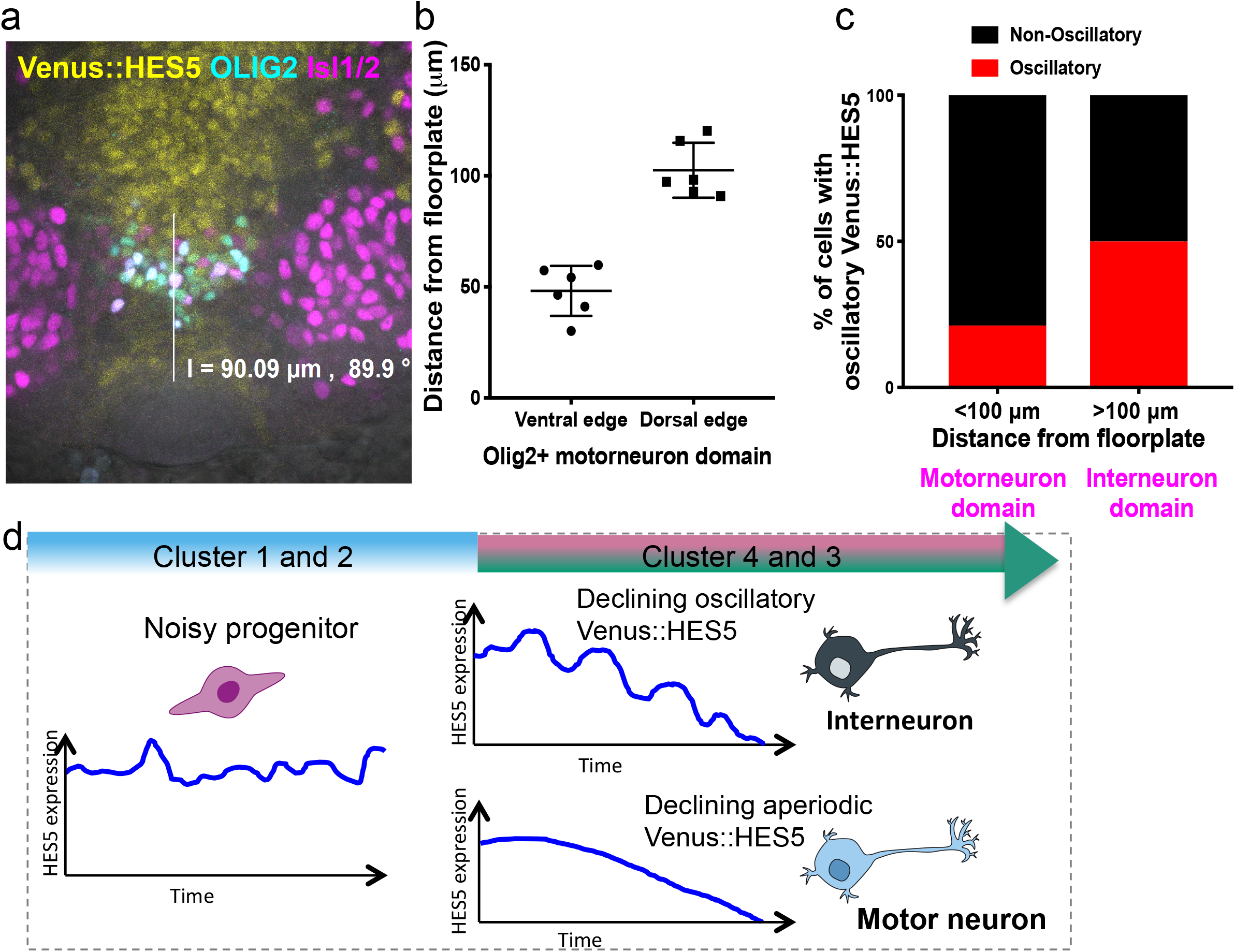
HES5 oscillations during differentiation correlate with cell fate. **a**) Immunofluorescence of E10.5 Venus::HES5 transverse slice of spinal cord ex-vivo. Endogenous Venus::HES5 signal, OLIG2 – motor neuron progenitors, Isl1/2 - mature motor neurons. Measurement bar from floorplate to dorsal edge of OLIG2 domain 90μm. **b**) Distance of ventral and dorsal edge of the motor neuron domain measured from the floorplate in ex-vivo spinal cord slice cultures. Bars indicate mean and error bars indicate SD n=6 slices. **c**) Percentage of single cells in cluster 3&4 classified as having oscillatory Venus::HES5 protein expression <100μm and >100 μm from floorplate. All cells (n=121) and experiments (n=3) combined. Fisher’s exact test performed on cell numbers in each category, p=0.004. **d**) Model of Venus::HES5 expression dynamics through cell-state transition from neural progenitor cell to neuron. Neural progenitors have dynamic, noisy and aperiodic fluctuations in Venus::HES5 protein expression. As cells transition towards neurons they have a long-term decreasing trend in Venus::HES5 and are more likely to show short-term oscillatory dynamics. Cells declining with oscillatory Venus::HES5 correlate with interneuron fate, whereas declining non-oscillatory cells correlate with motor neuron fate. Source data are provided in a Source Data file.

## Discussion

We have investigated how individual Sox1+ NPCs and their progeny make cell state transitions. Our main findings are twofold: firstly, oscillatory expression of HES5 is observed in NPCs in the tissue environment but occurs more frequently and with higher fold change in cells that are transitioning towards a differentiated interneuron state (Fig. 8d). Secondly, cell-to-cell heterogeneity in HES5 in tissue is a composite of long term dynamics (decline in expression) and short term dynamics (fluctuations in a short time scale).

Our findings support the view that changes in expression dynamics correlate with transitions in cell state^9^. However, contrary to expectations^13^ we observe both oscillatory and non-oscillatory HES5 dynamics within 2 defined sub-states; noisy HES5 dynamics in the proliferative progenitors and HES5 oscillatory dynamics being more likely in the cells differentiating towards interneurons. Our findings extend previous data, as the HES5 amplitude, period and dynamic behavior in tissue with statistical and computational tools have not previously described.

The HES5 oscillator operates around a high mean with low peak-to-trough amplitude in dividing progenitor cells (clusters 1 and 2). The small differences in peak and trough levels may be difficult to differentially decode by downstream genes. Most likely, these oscillations are a by-product of an active negative feedback loop that is required for maintaining the HES5 level around a high mean, thus repressing pro-neural genes in most apical progenitors. By contrast, oscillations in the transition to differentiation are coupled with an overall declining trend, and thus generate larger fold differences, which may be easier for downstream targets to decode. This is analogous to a ball bouncing down steps and undergoing greater instantaneous height drops (oscillatory expression) than a ball rolling down a ramp (aperiodic expression). Since HES proteins are transcriptional repressors for pro-neural genes, such as *Neurog2* and *Atoh1*^21,51^ we predict that the larger fold-changes generated by oscillatory decline in HES5 induces an oscillatory onset of downstream proneural genes^11,35^. We argue that coupling HES5 oscillations with a declining trend is an ingenious biological way for the cells to be able to decode what is normally a very shallow HES5 oscillator and importantly, to couple it with the process of differentiation. While it is known that HES5 in motor neurons is downregulated by OLIG2^7^ it is not clear what causes the decline in interneurons.

Our mathematical modelling identified that HES5 oscillations are enabled by stochastic amplification^48^ and that the HES5 auto-repression network operates near a bifurcation boundary. Consequently individual cells can switch between aperiodic and periodic expression stochastically and through regulated parameter changes. Instead of considering oscillatory versus non-oscillatory cells as stable distinct subpopulations, we propose that these are readily interconverting states with considerable plasticity. The model predicts that the transition towards oscillatory behaviour is most likely regulated through changes in the Hill coefficient, repression threshold or protein degradation. Future development of this model will capture remaining features of the observed dynamics such as the down-regulation observed during the differentiation process, as observed in cells of cluster 3 and 4, as well as other gene regulatory interactions and multi-cellular interactions.

In our modeling, we have included the effects of intrinsic stochastic noise, since this does not introduce further model parameters and is associated with any rate process. Phillips et al.^52^ suggested that low HES1 molecule number leads to stochastic oscillations of HES1 through a finite number effect. By contrast, HES5 molecule number is not low, (approximately 30-55k molecules per nucleus for HES5 versus while 2-3K per nucleus for HES1^30^). Thus other sources of noise may need to be considered perhaps stochastic activation of Notch cell-cell signalling in the densely packed tissue or cell division^53^ and the cell cycle. Noise and stochasticity are often considered undesirable yet they may also benefit decision making processes^30^. Here the benefit of noise may be to prime HES5 expression such that it is poised to become oscillatory.

The increase in cells with oscillatory HES5 in dissociated cells versus cells in tissue, is in agreement with previous suggestions of a cell-autonomous *Hes* oscillator which can be tuned by external signals^24,52^. The lack of statistical difference between the number of oscillating cells from homozygous and heterozygous Venus::HES5 animals suggests, that in contrast to the stochastic transcriptional bursting^54^, negative feedback generated oscillations can be somewhat synchronous between 2 alleles.

Other genes in the Notch-Delta network such as *Hes1, Dll1* and *Neurog2* have been shown to oscillate in NPCs. The relative timing of pulses of different genes may regulate cellular behaviour as common target genes may respond differently to inphase or out-of-phase input pulses. Indeed, the relative phase of the Notch and Wnt signalling oscillations in somitogenesis have been proposed to control cellular differentiation^55^. Imaging protein expression dynamics of multiple factors in the same cell during cell fate decisions would help to reveal the relative timing of multiplexed oscillatory gene expression.

The second main contribution of this paper is to increase the depth of our understanding of the degree and origin of cellular heterogeneity in gene expression in a tissue environment. We conclude that HES5 expression in the spinal cord is not an ergodic system since tissue level variability cannot be explained from short-term single cell variability but through a combination of cell sub-states co-existing in the tissue (which can be resolved spatially and dynamically) and transitions between these sub-states. A progenitor zone close to the ventricle (<50μm) shows maximum heterogeneity in cell-states, as all 4 dynamic expression clusters are equally represented in this zone, but minimum cell-to-cell heterogeneity in HES5 expression levels. By contrast, in the progenitor zone further from the ventricle there is minimum heterogeneity in cell-states, as it occupied mainly by cells in clusters 3 and 4, and maximum cell-to-cell heterogeneity in HES5 expression levels, approaching a 10-fold range in HES5. Furthermore, single cells undergoing differentiation start to down-regulate Venus::HES5 at any point between 20-50μms away from the ventricle indicating that cells can make the cell fate decision at any point along the apical-basal dimension of the progenitor zone. Though we could not resolve the differences between clusters 1 and 2, and clusters 3 and 4, our findings contrast with the schematic view that cell fate is controlled deterministically at global tissue level through signalling gradients. Together with the finding that more cells show oscillations in a dissociated culture, we suggest that NPCs make stochastic fate decisions through a complex and yet unresolved integration between global and local cell-cell signalling.

Our findings highlight the importance of integrating gene expression dynamics with spatio-temporal cell behavior to understand cell state transitions in real time in a multicellular tissue.

## Methods

### Animal models

Animal (*Mus musculus*) experiments were performed under UK Home Office project licenses (PPL70/8858) within the conditions of the Animal (Scientific Procedures) Act 1986. Animals were only handled by personal license holders. Venus::HES5 knock-in mice (ICR.Cg-Hes5<tm1(venus)Imayo>)^9^ were obtained from Riken Biological Resource Centre, Japan and mated with CD-1 mice for 1 generation before being maintained as an in-bred homozygous line. In these mice the mVenus fluorescent protein is fused to the N-terminus of endogenous HES5. R26R-H2B::mCherry mice^56^ were obtained as frozen embryos from Riken Centre for Life Science Technologies, Japan and C57Bl6 mice were used as surrogates. Sox1Cre:ERT2 mice (Sox1tm3(cre/ERT2)Vep^57^ were obtained from James Briscoe with the permission of Robin Lovell-Badge. Sox1Cre:ERT2 (NIMR:Parkes background) and R26R-H2B::mCherry (C57Bl6 background) were crossed to generate a double transgenic line (mixed background) homozygous for R26R-H2B::mCherry and heterozygous for Sox1Cre:ERT2.

### Embryo slicing

Homozygous Venus::HES5 knock-in females were mated with R26R-H2B::mCherry Sox1Cre:ERT2 males and E0.5 was considered as midday on the day a plug was detected. Intra-peritoneal injection of pregnant females with 2.5 mg Tamoxifen (Sigma) was performed 18 hours prior to embryo dissection. Whole embryos were screened for H2B::mCherry expression using Fluar 10x/0.5 objective on a Zeiss LSM880 confocal microscope and the trunks of positive embryos were embedded in 4% low-gelling temperature agarose (Sigma) containing 5mg/ml glucose (Sigma). 200μm transverse slices of the trunk around the forelimb region were obtained with the Leica VT1000S vibratome and released from the agarose. Embryo and slice manipulation was performed in phenol-red free L-15 media (ThermoFisher Scientific) on ice and the vibratome slicing was performed in chilled 1xPBS (ThermoFisher Scientific).

### Fluorescence Correlation Spectroscopy

E10.5 transverse spinal cord slices heterozygous or homozygous for Venus::HES5 were stained on ice for 1.5 hours with 50μM Draq5 (ThermoFisher Scientific) diluted in phenol-red free L-15 (ThermoFisher Scientific) media. Fluorescence Correlation Spectroscopy (FCS) experiments and snapshot images of whole spinal cord were carried out using a Zeiss LSM880 microscope with a C-Apochromat 40x 1.2 NA water objective on slices placed directly on a glass-bottomed dish (Greiner BioOne) kept at 37°C and 5%CO_2_. FCS signals were collected inside single nuclei in either the ventral region alone or both dorsal and ventral regions for tissue experiments. Venus (EYFP) fluorescence was excited with 514 nm laser light and emission collected between 517 and 570nm. Data from individual cell nuclei was collected using 5 x 2 s runs at 0.15 to 0.3% laser power which gave <10% bleaching and a suitable count rate ~1 kHZ counts per molecule (CPM). To obtain molecule number, autocorrelation curves were fit to a two-component diffusion model with triplet state using the Levenberg-Marquardt algorithm in MATLAB optimization toolbox with initial conditions assuming a ‘fast’ diffusion component 10x faster than the ‘slow’ component^58^. Measurements collected from cells exhibiting large spikes/drops in count rate or with low CPM (<0.5 kHz), high triplet state (>50%), or high bleaching (>10%) were excluded from the final results. Number and brightness analysis of the count rate^59^ showed a high correlation with molecule number obtained from autocorrelation curve fitting. The effective confocal volume had been previously determined to be 0.57fL ± 11 fL (mean with S.D.) using Rhodamine 6G with known diffusion constant of 400 μm^2^ s^−1^ allowing conversion from molecule number to concentration^60^. Single-cell data of number of molecules in the cell nucleus was obtained by adjusting concentration to the average volumetric ratio between nuclear volume and confocal volume. Mean nuclear volume of 523 fL was estimated using H2BmCherry intensity and 3D reconstruction from z-stack images in Imaris (Bitplane).

### Generating a quantitative expression map

Individual Draq5+ nuclei in a tile-scan image of a transverse slice of the whole E10.5 spinal cord were manually segmented as ellipses using ImageJ and background Venus::HES5 fluorescence (measured via an ROI drawn outside of the cells) was subtracted. A quantile-quantile plot was generated for the distribution of nuclear Venus::HES5 intensities from manual segmentation of a single image and the distribution of nuclear Venus::HES5 concentrations from FCS of cells throughout the E10.5 spinal cord from multiple slices and experiments. Linear regression was used to generate a calibration curve between Venus::HES5 intensity and Venus::HES5 concentration over the middle 90% of the range. The gradient of the line was used as a scaling factor and applied to the pixel intensity values in the segmented image to transform intensity to concentration.

### Analysis of variability in Venus::HES5 in snapshot images

The centroids of the manually segmented cells from a quantitative expression map were used to measure distance from the ventricle and perpendicular to the D/V axis. Neighbours were ranked based on distance from the centroid of the cell of interest and the nearest neighbours were classified as the cells in the first rank (Supplementary Fig. 1n). Coefficient of variation of Venus::HES5 intensity was measured by manual segmentation of Draq-5 stained transverse slices of whole E10.5 spinal cord in ImageJ.

### Embryo slice culture and live imaging

E10.5 spinal cord slices for live timelapse microscopy were placed on a 12mm Millicell cell culture insert (MerckMillipore) in a 35mm glass-bottomed dish (Greiner BioOne) incubated at 37°C and 5%CO_2_. The legs of the cell culture insert were sanded down to decrease the distance from the glass to the tissue. Primary neural stem cells dissociated from E10.5-E11.5 Venus::HES5 spinal cords were maintained as a line for up to 10 passages and plated in 35mm glass-bottomed dish (Greiner BioOne) for live imaging. 1.5mls of DMEM F-12 (ThermoFisher Scientific) media containing 4.5mg/ml glucose, 1x MEM non-essential amino acids (ThermoFisher Scientific), 120ug/ml Bovine Album Fraction V (ThermoFisher Scientific), 55μM 2-mercaptoethanol, 1x GlutaMAX (ThermoFisher Scientific), 0.5x B27 and 0.5x N2 was added. Movies were acquired using Zeiss LSM880 microscope and GaAsP detectors. For slice imaging a Plan-Apochromat 20x 0.8 NA objective with a pinhole of 5AU was used. 10 z-sections with 7.5 μm interval were acquired every 15 mins for 18hrs. DMSO (Sigma) or 2μM DBZ (Tocris) was added to media immediately before imaging. For imaging dissociated cells a Fluar 40x 1.3 NA objective with a pinhole of 6.5AU was used. 6 z-sections with 3.7 μm interval were acquired every 10 mins for 24-48hrs.

### Image analysis and cell tracking

Briefly, single cells were tracked using the H2B::mCherry channel. Background fluorescence as measured via an ROI drawn on a non-Venus::HES5 expressing region on the tissue was subtracted prior to analysing time-lapse intensity data. For cells in the ex-vivo slices single-cell Venus and mCherry expression were normalised to the whole tissue mean for the relevant channel to account for any possible photobleaching. For hierarchical clustering single-cell Venus::HES5 expression from 12-hour tracks was standardized by subtracting the mean and dividing by the standard deviation of the single-cell signal. Time 0 refers to the start of the tracking and not necessarily the start of the movie.

Single neural progenitor cells in E10.5 spinal cord slices were tracked in Imaris on the H2BmCherry channel using the ‘Spots’ and ‘Track over time’ function. Spot detection algorithm used background subtraction and tracking used the Brownian motion algorithm. All tracks were manually curated to ensure accurate single-cell tracking. A reference frame was applied to the movie along the dorso-ventral and apico-basal axes of the spinal cord to allow the distance from the ventricle to be calculated. To account for any photobleaching and allow comparison of intensities between movies the mean intensity of mCherry and Venus in each spot was normalised to the mean intensity of mCherry or Venus in the whole tissue. The whole tissue volume was tracked using the ‘Surfaces’ and ‘Track over time’ function.

There was no correlation in Venus::HES5 and H2BmCherry expression suggesting the Venus::HES5 dynamics were not a result of global changes in transcription or translation in the cell or microscope anomalies (Supplementary Fig. 3a-d). We also investigated the relationship between Venus::HES5 and z-position in the tissue (Supplementary Fig. 3e-h). As expected from imaging through tissue there was a small negative correlation (r = −0.24) between Venus::HES5 intensity and z-position when all cells and time-points were plotted (Supplementary Fig. 3f). However the range of z-positions in a single cell 12-hour track was rarely greater than 25μm, therefore it is unlikely the fluctuations and oscillations in Venus::HES5 are a result in changes in z-position (Supplementary Fig. 3g). Further at the single-cell level there is no difference in the correlation coefficient between z-position and Venus::HES5 intensity when comparing oscillatory and non-oscillatory cells (Supplementary Fig. 3h).

To compare levels of Venus::HES5 expression between nuclei and movies the effect of increased light scattering with increasing depth of tissue was corrected to the initial z-position of the nuclei in the tissue. This was only performed when a comparison of absolute levels within the movies was required. For each movie, a plot of single cell z-depth in the tissue vs single cell Venus::HES5 intensity was performed for all data points (similar to Supplementary Fig. 3f). Linear regression using least squares was performed to find the slope and y-intercept of the relationship between z-depth and Venus::HES5 intensity. For the initial time point the Venus::HES5 intensity was then corrected as if the cell was at z-position 0 by multiplying the slope of the z vs intensity relationship with the initial z-position and adding this value to the observed Venus::HES5 intensity. This value was then added to the Venus::HES5 intensity at all subsequent timepoints.

### Hierarchical clustering

Prior to analysis, timeseries of single cell Venus::HES5 expression were normalised to tissue mean to account for bleaching per independent experiment and in addition standardised (z-score calculation) by subtracting the mean of the timeseries from each timepoint and dividing by the standard deviation of the timeseries. Standardising the data enables clustering on relative expression changes rather than absolute expression levels. Cells were aligned to all start at time 0, which refers to the start of the tracking rather than the start of the movie. Standardized single cell timeseries were then subject to hierarchical clustering using Euclidean distance and Ward’s linkage in RStudio (R Project). Experiments were clustered separately and each clustergram independently identified 4 clusters per experiment. The elbow method to look at the variance explained as a function of number of clusters (nbclust package, R), suggested 4-6 clusters as the optimal cluster number however 5 and 6 clusters were not favoured by silhouette method (nbclust package, R) so we chose 4 clusters. Cluster relationships varied between experiments thus for annotation between experiments corresponding clusters labels were determined by 1. observation of mean Venus::HES5 expression over time per cluster and 2. calculating average single-cell coefficient of variation (COV) in Venus::HES5 over time for each cluster and comparing to results of clustering experiment 1 (Supplementary Fig. 4b). Thus, four clusters with the same mean Venus::HES5 expression dynamics and COV profile are reproducibly identified in each experiment. For DBZ-treated cells, data could not be corrected for photobleaching since Venus::HES5 downregulation is induced at tissue level causing a significant drop in tissue mean and masking effects from bleaching. Prior to analysis both DMSO and DBZ timeseries were standardised by z-scoring. To enable comparison between DBZ-treated and negative control DMSO-treated cells, experimental data from both treatment conditions were clustered together (Fig. 4d) as well as clustering DMSO independently of DBZ (Supplementary Fig. 6) yielding similar cluster profiles to untreated cells (Supplementary Fig. 6f,g).

### Estimation of cell-cycle phases

Cell cycle phases were inferred based on position and trajectories of single nuclei over time. Nuclei were classified as in G1 if moving basally, following division, or if they maintained a basal position for multiple hours. Nuclei were classified as in S if they were in a basal position before moving apically and dividing. Similarly nuclei were classified as in G2 if they moved apically and divided. Cells were classified as undergoing mitosis if the H2B::mCherry signal was observed to duplicate. Multiple cell-cycle phases were attributed to each cell and all phases used to calculate a percentage profiles.

### Analysis of long-term trends in Venus::HES5 expression

For 4, 8, 12, 14.25 and 17.25 hour time windows the coefficient of variation (standard deviation/mean x100) of all the normalised Venus::HES5 intensity values for a single cell in the time window was calculated. The shoulder point of Venus::HES5 was defined as a turning point in the signal that lead to a decrease of greater than 50% of the signal.

### Immunofluorescent staining

Trunks of E10.5 embryos for cryo-sectioning were fixed in 4% PFA for 1 hour at 4°C, followed by 3 quick washes with 1xPBS and 1 longer wash for 1 hour at 4°C. Embryos were equilibrated overnight in 30% sucrose (Sigma) at 4°C before mounting in Tissue-Tek OCT (Sakura) in cryomoulds and freezing at −80°C. 12μm sections were cut on Leica CM3050S cryostat. E10.5 spinal cord slices cultured on Millicell inserts were fixed in 4% PFA for 4 hours. For staining, tissue and sections were washed in PBS followed by permeabilisation in PBS 0.2% Triton X-100 (Sigma) and blocking with PBS 0.05% Tween20 (Sigma) + 5% BSA (Sigma). Primary and secondary antibodies were diluted in PBS 0.05% Tween20 + 5% BSA. Tissue was incubated with primary antibodies overnight at 4°C, then washed three times for 5–10 minutes in PBS 0.05% Tween20, incubated with secondary antibodies and DAPI (Sigma) for 4 hours at room temperature, and washed again three times in PBS-T. Sections were mounted using mowiol 4-88 (Sigma). Primary antibodies used were rabbit anti-SOX2 (ab97959, 1:200), mouse anti-NeuN (Merck MAB377, 1:100) mouse anti-NKX2.2 (74.5A5, Developmental Studies Hybridoma Bank, 1:10), mouse anti-PAX7 (Developmental Studies Hybridoma Bank, 1:10), rabbit anti-Olig2 (EMD Millipore AB9610, 1:200), mouse anti-Isl1/2 (Developmental Studies Hybridoma Bank, 1:100) and rabbit anti-β3-tubulin (Cell Signaling Technology, 5568S 1:200).

### Cell culture

Primary NS cells were isolated from dissected spinal cords of E10.5-11.5 embryos from Venus::HES5 knock-in mice and cultured in DMEM/F-12 containing 4.5mg/ml glucose, 1x MEM non-essential amino acids (ThermoFisher Scientific), 120ug/ml Bovine Album Fraction V (ThermoFisher Scientific), 55μM 2-mercaptoethanol, 1x GlutaMAX (ThermoFisher Scientific), 0.5x B27 and 0.5x N2. NS-E cells were a gift from Jennifer Nichols (Cambridge Stem Cell Institute, UK).

### Half-life experiments

Protein half-life was obtained by transfection of 3xFlag-HES5 and 3xFlag-Venus::HES5 in to NS-E cells with Lipofectamine 3000 (ThermoFisher Scientific) as per manufacturers’ instructions. 24 hours after transfection, cells were treated with 10μM cycloheximide (Sigma) and at 0, 15, 30, 60, 120, and 240 mins after treatment lysed with. Western blots were performed using 4-20% Tris-glycine acrylamide gels (NuSep), Whatman Protran nitrocellulose membrane (Sigma) and developed with Pierce ECL substrate (ThermoFisher Scientific). Antibodies used were anti-HES5 [EPR15578] (Abcam, ab194111) and anti-alpha-tubulin (clone DM1A Sigma T9026). RNA half-life experiments were obtained by 10μM actinomycin D (ThermoFisher, Scientific) treatment of primary heterozygous Venus::HES5 and primary wild-type spinal cord NS cells. Samples were taken at 0, 15, 30, 45, 60, 80, 100, 120 mins after treatment and RNA prepared using RNAeasy kit (Qiagen) with DNAse treatment as per manufacturers instructions. cDNA was prepared using Superscript III (Invitrogen) as per manufacturers’ instructions and qPCR for Venus, HES5 and GAPDH was performed with Taqman (ThermoFisher, Scientific, UK) gene expression assays.

### Statistical testing

Statistical tests were performed in GraphPad Prism 7. Data was tested for normality with D’Agostino-Pearson test. The relevant parametric or non-parametric test was then performed. If necessary outlier removal was performed using ROUT method (GraphPad). Coefficient of variation is defined as standard deviation (SD) over the mean.

Stacked bar plots and discrete scatter plots show mean or mean±SD where multiple independent experiments are analysed. Statistical significance between 2 datasets was tested with either Student t-test (parametric) or Mann-Whitney test (non-parametric). Statistical significance (p<0.05) for 2+ datasets was tested by Kruskall-Wallis with Dunn’s multiple comparison correction. All tests were 2-sided. Multiple comparison testing involved comparing all pairs of data columns. Correlations were analysed using Spearman rank correlation coefficient. Sample sizes, experiment numbers, p values<0.05 and correlation coefficients are reported in each figure legend.

### Detection of oscillations using Gaussian processes

We adapted the statistical approach developed by Phillips et al.^43^ to analyse timeseries of Venus::HES5 in single cells tracked in 3D fluorescence imaging. Data was de-trended to remove long term behaviour such as down-regulation and to recover the oscillatory signal with zero mean. We used maximum likelihood estimation to fit the de-trended data timeseries with two competing models: a fluctuating aperiodic one (null model) and an oscillatory one (alternative model). We used the log-likelihood ratio statistic to compare the likelihood of data being oscillatory or non-oscillatory and determined the oscillators based on a false discovery rate of 3% independently per experiment. The custom algorithms and routines were implemented and tested in MATLAB R2015a and are using the GPML toolbox^61^.

### Gaussian processes (GPs) background

GP inference is a probabilistic modelling technique that involves fitting a time-series signal *y*(*t*) in terms of a mean function, *m*(*t*) describing the moving average of signal with respect to time and a covariance function, *k*(*τ*) describing how the signal varies around the mean with respect to time

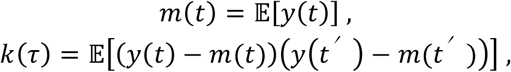

where the covariance function for this process varies with *τ* = |*t – t*′|, representing the time interval between any pair of time points (*t,t*′). The covariance function is typically represented by a parameterised function that encapsulates modelling assumptions. The GP provides a generative model for the data, i.e. for known mean and covariance functions synthetic data may be generated by sampling from the multivariate normal distribution 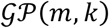 evaluated at the times where data are collected. The likelihood function, which is the probability of the observed data under the model, is exactly tractable making inference and parameter estimation possible.

### Detrending single cell expression timeseries

Single cell protein expression timeseries contain information about dynamics at long timescales (above 10-12hrs) as well as dynamics at short timescales (2-5hrs). To account for this, we first model the long-term behaviour (i.e. mean function) using a squared covariance function:

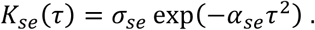

This allows us to determine the mean function 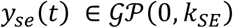 and extract the detrended dynamics by: *y_d_*(*t*) = *y*(*t*) − *y_se_*(*t*). We used a lengthscale *a_se_* corresponding to 10hrs to remove long term dynamics while preserving short periodicity dynamics. Next we modelled the detrended data with zero mean using GP and two competing covariance models each with characteristic parameters inferred from the data. These models are described in the following section.

### Oscillatory and Non-Oscillatory Covariance models

The detrended timeseries can be oscillatory or non-oscillatory (examples in Supplementary Fig. 8a and Supplementary Fig. 8b respectively). To account for this, two covariance models are used, namely:

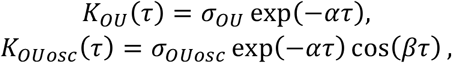

where *K_OU_* is a standard Ornstein-Uhlenbeck (OU) covariance function, which models *aperiodic* stochastic fluctuations, while *K_OUosc_* is a oscillatory OU covariance function, which models *periodic* stochastic timeseries. The parameters determining these models are:

- signal variance *σ_OU_, σ_OUosc_*; related to signal amplitude by 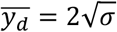;
- lengthscale *α* which represents the rate at which correlations between subsequent peaks decay over time (see also discussion in Prior Distribution on Lengthscale);
- frequency *β*; related to periodicity by *T* = 2*π*/*β*.

In the following section, we discuss how the probability of the observed data given the periodic model is affected by technical noise and we introduce a global calibration technique that accounts for this.

### Global Calibration of Technical Noise

The fluorescent detrended signal from an oscillatory cell contains periodic variations over time (modelled by the stochastic and periodic covariance model *K_OUosc_*) as well as additive technical noise: 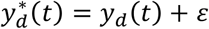, where *ε* denotes white noise of zero mean and variance *σ_n_*. The technical noise parameter *σ_n_* is unknown and needs to be estimated.

Phillips et al.^43^ use an experimental estimation of technical noise where signal collected from empty areas of the cell culture dish (background) is used to approximate signal detection noise in bioluminescence imaging. Here we use a fluorescence reporter which suffers from variability of noise from the detector (also present in bioluminescence) and auto-fluorescence produced by the cells thus showing an overall increased noise level in cells than background.

To account for this, we propose a different strategy to estimate technical noise directly from the data by observing the relationship between likelihood and signal-to-noise ratio, i.e. optimizing joint likelihood. The joint likelihood function describes the probability of observing all the data per experiment under a global model:

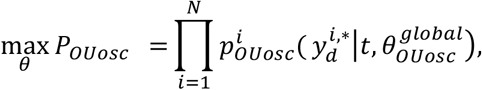

where 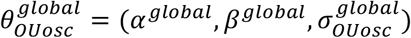 represents the global hyperparameters and 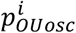 denote individual likelihood functions; and N is the total number of cells in each dataset. An equivalent and more convenient optimisation is to maximize the joint log-likelihood function:

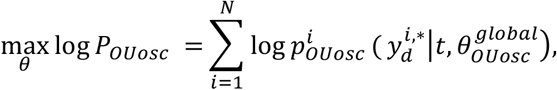

where the cell-specific log-likelihood functions are:

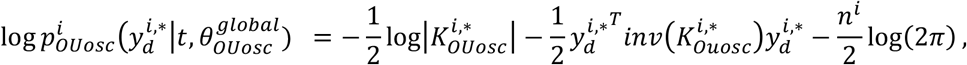

with *n^i^* representing the number of time-points in each trace and 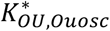 denoting the cell-specific OUosc covariance models with technical noise:

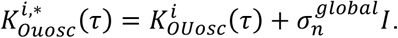

The global estimation of 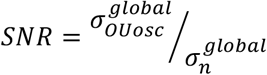 indicates that the joint log-likelihood function is low (indicating a poor model) when the signal is assumed to have high signal-to-noise and is highest (indicating an optimal model) for noise levels from 10-15% (Supplementary Fig. 8c). Since the joint log-likelihood levels are not improved for noise above 10%, we conclude that the characteristic level of noise at maximum joint log-likelihood for the 3D tissue data is approximately 10%.

For single cell parameter estimation (described in the next section), we calibrated the technical noise level expressed in fluorescent intensity units at the global level and re-estimated the remaining parameters for each cell. In this way we ensure that cells have the same (global) amount of technical noise variance and that the remaining variability comes from the true signal. This generates a fit that is robust and avoids over-fitting. In the following, we describe the parameter estimation procedure:

### Maximum Likelihood Estimation and Log-Likelihood Ratio

Using the global calibrated level of technical noise, we further optimise GP covariance model parameters for single cell timeseries. Since we do not know a priori if single cell traces are periodic or not, we estimate parameters for both OU and OUosc models by estimating the optimal hyperparameters 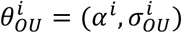 and 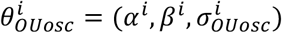 respectively. These parameters maximising the log-likelihood function:

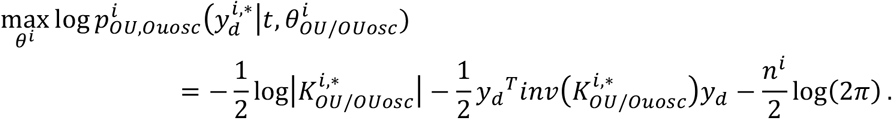

The likelihood value of the periodic model does not directly indicate if the data is oscillatory or non-oscillatory since OUosc can also fit non-oscillatory data (OUosc becomes OU for *β* → 0). However, a higher likelihood for OUosc compared to likelihood of OU indicates increased probability of the signal to be oscillatory. To quantify the probability for the single cell data to be oscillatory, we use the log-likelihood ratio statistic:

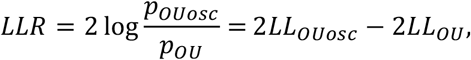

where 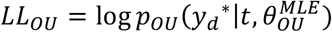 and 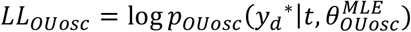. To account for effects caused by length of each trace *n^i^*, the following normalisation was used *LLR^i^* = *LLR^i^/n^i^*.

### Detection of Oscillators by False Discovery Rate (FDR)

In order to classify cells into oscillatory and non-oscillatory, we identify statistically significant LLR scores by comparing to LLR scores obtained from a null (non-oscillatory) distribution^62^ and controlling for the false discovery rate (FDR) per experiment. We obtain a synthetic null distribution using the generative OU (non-oscillatory) models that have been fitted to data. This allows generating a large number of synthetic traces that are non-oscillatory. To find statistically significant oscillators we set FDR to 3%. Controlling the FDR ensures that false positives are very unlikely to be considered oscillatory (less than 1 in 33 cells analysed) and an example from data is shown in Supplementary Fig. 8d. In combination with GP models used in this study, the FDR technique^63^ has been shown to outperform standard Lomb-Scargle periodogram (for detection of oscillations by frequency analysis) in terms of specificity and sensitivity^43^.

### Prior Distribution on Lengthscale

Lengthscale is a parameter of the covariance model that describes the rate at which subsequent peaks in the oscillatory signal become uncorrelated over time. In practice, we found that estimating lengthscale as a free parameter can lead to *α* → 0 for some of the cells (Supplementary Fig. 8e-panel 1 and insert). This possibility is unrealistic since it would imply correlations in the signal never decay over time. This vulnerability is likely to be caused by issues with the length of the data tracks which contain few samples (approximately 45 data points per cell acquired at 15min intervals) thus affecting the maximum likelihood technique.

To address this, we estimated lengthscale globally using the same technique described in Global calibration of technical noise section and used this to initialize the single cell parameter inference. In addition, we introduced a prior on the lengthscale that contains the global estimated value. The prior is defined as

SmoothBox1 (*SB*1) (existing in GPML toolbox) that has a sigmoidal expression around a lower bound, *l* and an upper bound, *L*:

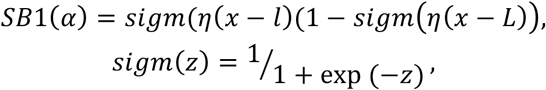

where *η* is a parameter that controls the shape, with higher values leading to a box appearance. The shape of the prior affects the posterior distribution of lengthscales that is estimated from the data. Although both a restrictive, box-shaped prior and relaxed, smooth prior help prevent *α* → 0 (Supplementary Fig. 8e-panels 2-3), the restrictive choice significantly alters the shape of the distribution at either bounds (Supplementary Fig. 8e-panel 2). By using a relaxed prior (Supplementary Fig. 8e-panel 3), we obtain a posterior distribution centered at the global estimate while still correcting for unwanted low values.

### Hilbert Reconstruction and Peak-to-Trough Fold Changes

We designed custom routines for instantaneous amplitude and phase reconstruction using the Hilbert transform and we used this to measure absolute peak-to-trough fold changes in signal over time. The Hilbert transform is used to reconstruct the instantaneous characteristics of the signal based on the analytic signal *z_d_*(*t*)^64^. The analytic signal consists of a real part identical to the data: *Re z_d_*(*t*) = *y_d_*(*t*); and an imaginary part representing the data with a *π*/2 phase shift, where phase shift denotes a difference in the peaks of two waves. It is generally assumed that, *Re z_d_*(*t*) is proportional to a cosine wave, thus *Im z_d_*(*t*) will be proportional to a sine wave or viceversa. The analytic signal relates to instantaneous amplitude *A* and phase *φ* by: *z_d_* = *A* exp(*iφ*). This leads to the following expression for amplitude and phase:

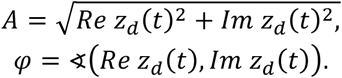

We applied the Hilbert transform implemented as *hilbert.m* in MATLAB on the fitted model obtained from analysing the detrended oscillatory data with *K_OUosc_*. Amplitude is detected in the detrended data and translated back to real intensity units (or relative units where appropriate) by summation with the long-term trend (Supplementary Fig. 8f). The reconstructed phase (Supplementary Fig. 8g) has a seesaw appearance that is characteristic of oscillatory data and indicative of a phase reset at the end of a complete period. Thus phase contains information on periodicity and we confirmed that periods estimated using GP agreed well with average reconstructed Hilbert periods (data not shown). In addition, the times at which the phase angle crosses the zero-level corresponds to peaks and troughs in the oscillatory signal. We used this property to identify peaks (ascending zero-crossing of phase) and troughs (descending zero-crossing of phase) in the real signal and generate absolute fold changes in amplitude by pairing a peak with the nearest trough and expressing the peak-to-trough intensity ratio for each pair. We report amplitude fold changes in the complete signal (containing the long-term trend and named peak:trough fold changes in figures and main text) either as maximum fold change or average fold change as indicated in figure legends.

### Stochastic model of HES5 expression dynamics

We modelled protein expression dynamics emerging from transcriptional autorepression and delay using an established mathematical model^46^. The model simulates changes of mRNA and protein number in a cell over time by considering the effects of transcription, translation, and degradation of protein and mRNA (Fig. 6a). The model includes effects of a transcriptional delay, representing that it takes a finite amount of time for mRNA to be produced and transported out of the nucleus for translation. The model further includes the effect of genetic auto-repression, i.e. we assume that high abundance of HES5 can inhibit its own transcription. In order to be able to describe both aperiodic and oscillatory dynamics we account for intrinsic stochasticity that is typically associated with rate-processes^44,49^. The model is implemented using delayed Chemical Langevin equations of the form^65,66^

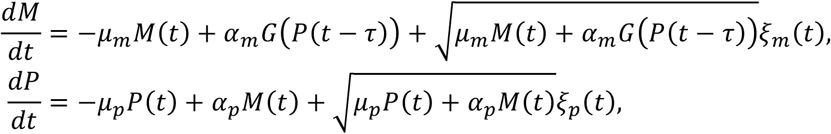

where *M*(*t*) denotes the number of mRNA molecules in one cell at time *t, P*(*t*) denotes the number of HES5 protein molecules, and *τ* represents transcriptional delay, i.e. the average time that is required for individual mRNA molecules to be transcribed and transported to the ribosome. The parameters *μ_m_, μ_p_, α_m_* and *α_p_* are rate constants denoting the rates of mRNA degradation, protein degradation, basal transcription in the absence of protein, and protein translation, respectively. The rate of transcription in the model is modulated in dependence of HES5 protein *P* at time *t* − *τ* by the Hill function^65^

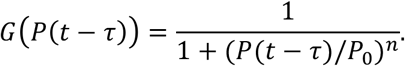

The variables *ξ_m_* and *ξ_p_* denote Gaussian white noise, which is characterised by delta-distributed autocorrelations^50^,

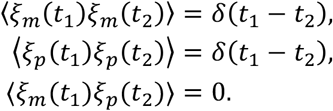

Note that Chemical Langevin equations allow non-integer values for the molecule numbers of mRNA and Protein.

### Numerical implementation of the model

We calculate summary statistics from the long-term trend of the model to enable model-data comparison. To investigate model behaviour at a unique combination of model parameters (in the following referred to as parameter point), we generate *N_p_* = 200 model traces at this parameter point and calculate averaged summary statistics from these traces. For each trace, the first *t*_eq_ = 1000 minutes are discarded and the remaining *t*_obs_ = 7500 minutes are used for the evaluation of summary statistics. The equilibration time *t*_eq_ is chosen such that that summary statistics do not depend on the initial condition. Initial conditions to evaluate the model at a given parameter point are *M*_in_ = 10 mRNA molecules, and *P*_in_ = *P*_0_ protein molecules. We impose that no mRNA transcription events were initiated at negative times by inhibiting transcription in the model for *t* < *τ*. Specifying model behaviour for negative times is necessary when evaluating delay differential equations. Chemical Langevin equations were implemented numerically using an Euler-Maruyama scheme and a time step of Δ*t* = 1 minute. All numerical simulation parameters are listed in Supplementary Table 1.

### Coherence and other summary statistics

To characterise the model behaviour at individual parameter points we collect the following summary statistics: (i) the *mean protein* expression level 〈*P*〉, (ii) the *relative standard deviation* of protein expression *σ_r,P_* = *σ_P_*/〈*P*〉, where *σ_P_* denotes the absolute standard deviation of protein expression, (iii) the *coherence* and (iv) the mean observed *period*. Means and standard deviations of expression (i-ii) are calculated across all traces at one parameter point and across all discretized timesteps in the observation window.

In order to calculate the oscillation coherence in Fig. 6d-g we use Fourier transforms of individual traces across the observation window, i.e

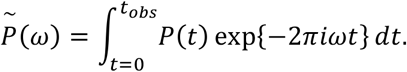

We then use the obtained Fourier transforms to calculate the power spectrum of the model at a parameter point, 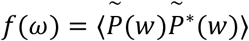. Here, the asterisk * denotes complex conjugation and the average is taken over all *N_p_* simulated traces at one parameter point. The discrete Fourier transform implemented in the Python numpy.fft package is used, introduced by Cooley and Tukey^67^. In order to minimise the influence of finite-size effects on the power spectrum we apply smoothing via a Savitzky-Golay filter^68^ with polynomial order three and a filtering window of 0.001/min, which corresponds to seven discrete frequency values. The coherence is then defined as *C* = *A*_max_/*A*_tot_^52,69^, where *A*_max_ is the area under the power spectrum in a 20-percent band centred at the frequency corresponding to the maximum of the power spectrum, and *A*_tot_ is the total area of the power spectrum. Typically, coherence is low for non-oscillatory cells and high for oscillatory cells. In Fig. 6h the linear noise approximation^44^ was used to calculate the power spectrum analytically before extracting coherence values, which reduces computational complexity. To calculate coherence values for the deterministic system^65^ (data shown in Fig. 6i) we used bifurcation analysis^70,71^ to identify whether oscillatory solutions exist at individual parameter points. If oscillatory solutions exist, the coherence at this parameter point is one, otherwise it is zero. In Fig. 6h,i the mean posterior predicted coherence values are plotted for varying degradation rates in the stochastic model and the deterministic model, respectively.

In order to extract period values from simulated data that can be compared to our experimental observations we applied the Hilbert transform technique (see also Hilbert Reconstruction) to simulated traces. At each parameter point, we generate one equilibrated trace of 12-hour duration, a similar observation window as the experimental data. Period values are identified as time differences between consecutive descending zero-crossings of the instantaneous phase, and the mean period across the measurement interval is recorded. In Fig. 6b the posterior predicted distribution of this mean period value is shown. The following section describes how Bayesian posterior predictions are generated.

### Parameter inference

In order to parameterise the model we use a combined approach of experimental parameter measurements and Approximate Bayesian Computation (ABC)^72,73^.

Specifically, we use experimentally measured values for the mRNA and protein degradation rates, corresponding to half-life values of 30 and 90 minutes, respectively (Supplementary Fig. 1). For the remaining, unknown, model parameters, we apply ABC, which is a standard method to infer parameters of non-linear stochastic models. ABC has the benefit of providing probability distributions for parameters, rather than point-estimates which in turn enables the estimation of parameter uncertainty, i.e. the uncertainty on the model parameters given the observed data.

We use ABC to identify parameter combinations that can explain key aspects of traces in clusters one and two. Specifically, we require the mean HES5 expression level to fall within the experimentally observed range of 55000 to 65000 protein molecules per cell (Supplementary Fig. 10a), and for which the standard deviation of modelled traces lies above five percent (Supplementary Fig. 10b). We analyse model predictions in Fig. 6 by investigating Bayesian posterior predictions, i.e. the distribution of model predictions given the posterior distribution of model parameters.

When investigating the impact of parameter changes on model predictions in Fig. 6j, we define a model prediction at a given parameter combination as aperiodic if (i) the predicted oscillation coherence is below 0.1, or if (ii) the predicted period, as identified by a peak in the power spectrum, is longer than 10 hours. We identify the posterior parameter samples fulfilling this condition and record their total number as *N_steady_*. Starting from these *N_steady_* aperiodic parameter combinations we change individual model parameters by 50% and count the number of parameter combinations *N_osc_* for which an individual parameter change leads to (iii) predicted coherence values above 0.1, and (iv) period values below 5 hours. The likelihood of an individual parameter change to induce oscillations in agreement with (i)-(iv) is then defined *L_osc_ = N_osc_/N_steady_*. We normalise the values of *L_osc_* to sum to a total probability of one in Fig. 6j. In order to ensure an accurate estimate of the likelihood values we increased the number of prior samples *N_tot_* to 2,000,000 in Fig. 6j, which ensured a sufficiently large *N_steady_*, with *N_steady_* = 20872.

### Background on Bayesian inference

In Bayesian statistics, the joint probability distribution *p*(**Θ,*D***) of a parameter vector (parameter point) **Θ** and observed data vector ***D*** is used to calculate the posterior distribution *p*(**Θ|*D***), the probability distribution of the parameters given the data. The calculation of the posterior is achieved by applying Bayes’ rule

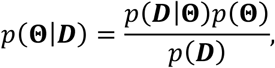

where *p*(***D*|Θ**) represents the probability of ***D*** given **Θ**, and is usually referred to as the likelihood, and *p*(***D***), the probability of observing the data, is the marginal likelihood. The likelihood of the parameter *p*(**Θ**) is referred to as the prior.

ABC can be used to estimate parameters of complex models for which the likelihood is not analytically tractable. Here, we apply rejection-based ABC^72^, which approximates the posterior through random sampling of parameters from the prior distribution and evaluating the model for each sample. Summary statistics are used in order to compare the model with data. This is necessary since the modelled data is high-dimensional: in our case it comprises the protein and mRNA numbers at each simulated timepoint in *N_p_* = 200 simulated cells per parameter point. As detailed above, we use the mean protein expression and the standard deviation of protein expression as summary statistics for fitting. In rejection-based ABC, samples for which the chosen summary statistics fall within an experimentally observed range are accepted as samples of the posterior distribution, otherwise they are rejected. Here, we generate *N*_tot_ = 200,000 samples from the prior distribution, providing us with 4,901 samples of the posterior distribution. The choice of prior distribution is discussed in the following section.

### Prior information on parameters

When applying ABC it is necessary to define a prior probability *p*(**Θ**)^73^. This prior is typically used to restrict inferred parameter combinations to biophysically realistic values, by imposing uniform and uncorrelated distributions on each parameter. We follow this approach by considering biophysical principles and literature values to define marginal prior distributions on each parameter as detailed below and summarised in Supplementary Table 2.

For the *basal transcription rate α_m_*, we assume a logarithmically-uniform prior, representing that we do not initially know the order of magnitude of this parameter. We consider a maximal, biophysically possible transcription rate of 60 transcripts per minute, which has previously been estimated^74,75^. We choose a lower bound of 0.1 transcripts per minute. This lower bound is chosen manually such that the prior bounds lie outside the support of the posterior. An example experimental estimate of a gene transcription rate is two transcripts per minute^76^. A genome-wide quantification of transcription rate estimates in mouse fibroblast cells revealed a distribution of transcription rates between 10^−3^ and eight transcripts per minute^77^.

For the *translation rate α_p_*, which represents the number of protein molecules generated per mRNA molecule per time interval, we use a logarithmically-uniform prior, similar to the prior in the basal transcription rate. We consider biophysical principles to impose an upper boundary on the translation rate as follows: ribosomes translate individual mRNA molecules with a speed of approximately six codons per second^78–80^. The footprint of an individual ribosome is approximately ten codons^81^. Thus individual ribosomes require approximately 1.6 seconds in order to free the translation start site, which limits the maximal possible translation rate to approximately 40 translation events per mRNA molecule per minute. We hence use 40/min as upper value for our prior on the translation rate, and the lower bound is chosen to be 0.5/min, which is chosen sufficiently small to lie on the outside of the support of the posterior. Genome-wide quantification of translation rate estimates included values of 20 translation events per mRNA per minute and above^77^.

The *repression threshold P*_0_ represents the amount of protein required to reduce HES5 transcription by half. For the repression threshold we use a uniform prior covering the range 0-120000. The upper bound is chosen as twice the value of the experimentally observed number of protein molecules per cell (~60,000), which is sufficiently large to include parameter regimes corresponding to no genetic autorepression.

The transcriptional delay *τ* corresponds to the time required to transcribe RNA and move it out of the nucleus for translation. Previous estimates of this parameter for a variety of genes varied between five and 40 minutes^82^, and typically assumed values are around 20-30 minutes^52,65^. Based on these values we used a uniformly distributed prior of five to 40 minutes.

The Hill coefficient *n* describes the steepness of the auto-repression response. Previously used values range between two and five^44,52,65,83^. Here, we use a uniform prior in the range of two to six. Importantly, we do not consider values above *n* = 6 since (i) the change in slope of *G* decreases for increasing *n*, and (ii) high values of *n* correspond to steep, step-like, response curves, which are unrealistic.

### Prediction of mean and variance correlation

In order to estimate how mean and variance inter-depend in our model we obtained Bayesian posterior predictions for mean HES5 expression and for its variance after changing the repression threshold (P_0_), which we varied from 10% to 200% of its original value in 10% intervals. For each relative change in the repression threshold these Bayesian posterior predictions are probability distributions analogous to those in Figure 6b and c. We then extracted the mean of the predicted HES5 levels, the mean variance of expression, and the standard deviation of the predicted variance for each relative change in repression threshold. In Supplementary Figure 11d we plotted the mean predicted level of HES5 against the mean variance obtained in this way, and we used the standard deviation at each relative value of the repression threshold as an estimate of the confidence interval of this prediction. As expected, the mean and variance of HES5 expression are positively correlated. The parameter variation of the repression threshold was conducted on 4901 posterior samples.

## Data availability

Single-cell Venus::HES5, H2B::mCherry intensity and positional information from ex-vivo movies are available in the source data and on request from the corresponding authors. The source data underlying Figs 1–5,7–8 and Supplementary Figs 1-9,11-12 are provided as a Source Data file. Simulations, data and code to generate Figs 6 and Supplementary Fig 10 are available online under https://github.com/kursawe/hesdynamics The raw movie files are available through figshare DOI 10.6084/m9.figshare.8005652.

## Code availability

Data fitting for detections of oscillations has been implemented in Matlab R2015a using the GPML toolbox (Rassmussen and Hannes 2010) and custom designed routines available at http://gaussianprocess.org/gpml/code/matlab/doc/. Code for stochastic model of transcriptional auto-repression and Bayesian inference are available online under https://github.com/kursawe/hesdynamics. Matlab custom designed routines for analysis of FCS available on request.

## Acknowledgements

We are grateful to Dr Raman Das and Dr Alexander Auhlela for help establishing slice culture methods and Drs. Ximena Soto and James Briscoe for advice and discussions. The authors would also like to thank the Biological Services Facility and the Bioimaging Facilities of the University of Manchester for technical support. This work was supported by a Sir Henry Wellcome Fellowship to CM (103986/Z/14/Z). VB and JK were supported by a Wellcome Trust Senior Research Fellowship to NP (090868/Z/09/Z). MR was supported by a Wellcome Trust Investigator Award (204832/B/16/Z). JB and DS were funded by MRC grants MR/K015885/1 and MR/M008908/1. The funders had no role in study design, data collection and analysis, decision to publish, or preparation of the manuscript.

## Author Contributions

CM and NP conceived and designed the experimental study. CM performed half-life experiments, acquired FCS data, acquired and analysed snapshot spinal cord slice images, acquired, tracked and analysed live spinal cord slice and dissociated cell imaging movies, performed cluster analysis and cell positional/migration analysis, interpreted data and wrote the paper.

VB developed method to detect oscillations in noisy timeseries data and their period, amplitude and fold-changes, analysed expression dynamics in single-cell data and wrote code to identify cell neighbours and extract positional information from singlecell tracking.

JB wrote custom code to analyse FCS data by auto-correlation with model fit and number and brightness, optimized settings for FCS in tissue environment and aided acquisition and performed Q-Q analysis to generate quantitative expression map.

JK designed efficient implementation of stochastic and deterministic HES5 models, planned and performed Bayesian inference to parameterize both models, analysed both models and performed bifurcation analysis.

BY performed immunohistochemical staining for D/V progenitor domain markers to map Venus::HES5 expression domains.

DS assisted with optimization of settings for FCS in tissue environment and imaging of slice cultures.

CMS supervised and assisted analysis and interpretation of FCS data.

TG supervised and assisted analysis and interpretation of HES5 model.

MR supervised and assisted development of method to detect oscillations in noisy timeseries data.

NP supervised and directed the work, interpretated data and co-wrote the paper with CM, VB and JK with input from JB and DS.

