## Supplementary Information for "Quantitative single-cell live imaging links HES5 dynamics with cell-state and fate in murine neurogenesis"

Manning et al.

**Supplementary Figure 1. Characterisation of Venus::HES5 in mouse E10.5 spinal cord**

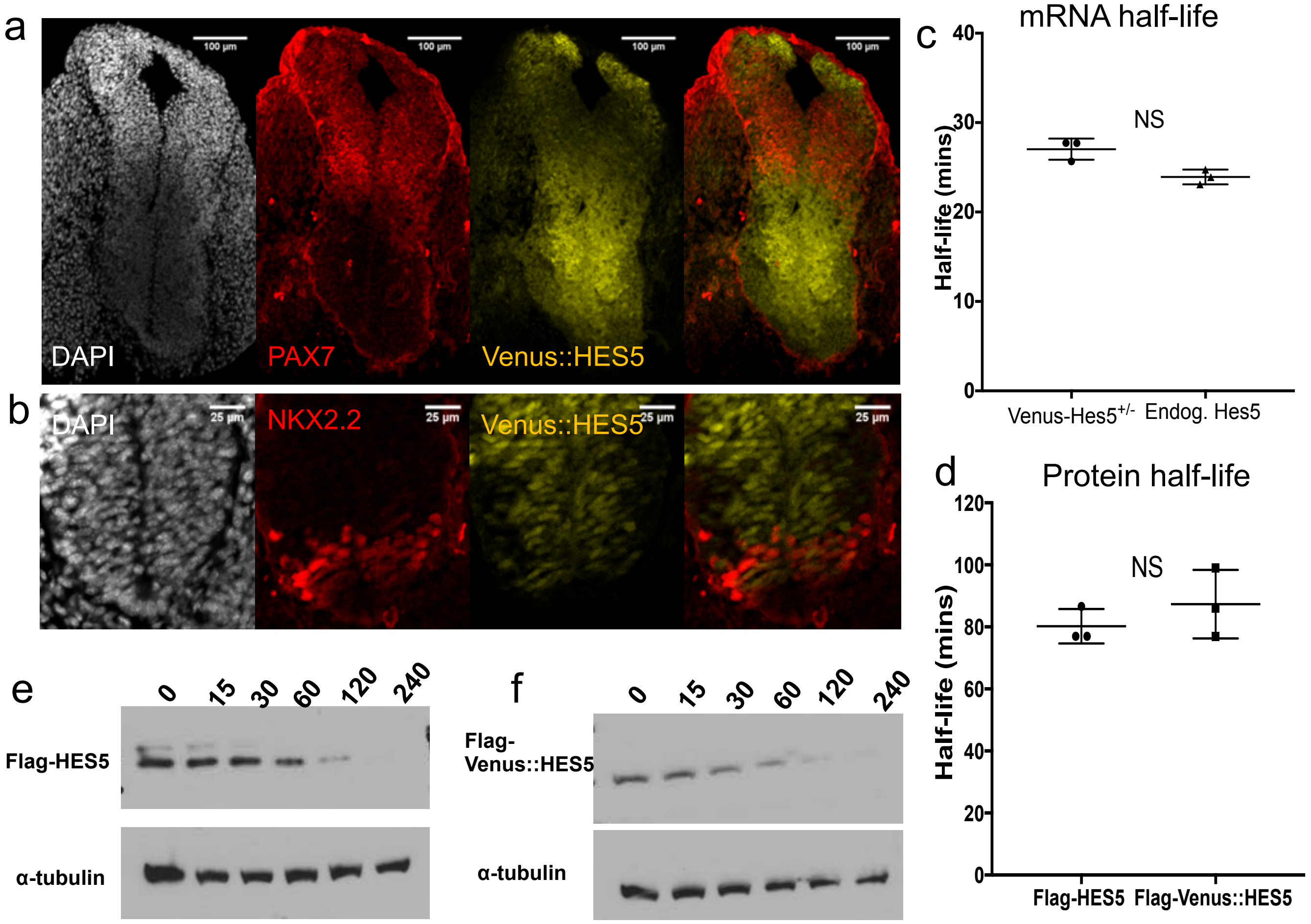


**Supplementary Figure 1. Characterisation of Venus::HES5 in mouse E10.5 spinal cord.** Immunofluorescence of frozen E10.5 Venus::HES5 knock-in mouse transverse section for **a**) PAX7, scale bar 100µm and **b**) NKX2.2 and endogenous Venus::HES5 signal, scale bar 25µm. The ventral Venus::HES5 domain is bounded dorsally by PAX7, which stops at a dorsal interneuron domain, dp6 and on the ventral side by NKX2.2, a ventral motoneuron (p3) marker. Thus, at E10.5, the ventral Venus::HES5 domain encompasses mainly progenitors of ventral interneurons (p0/p2) and of some ventral motorneurons (p2/pMN). **c**) mRNA half-life of *Venus::Hes5* or *Hes5* in primary neural stem cells from heterozygous Venus::HES5 E10.5 spinal cord embryos. n=3 experiments. **d**) Half-life of Flag-Venus::HES5 or Flag-HES5 protein measured in vitro. n=3 experiments. NS no significant difference in student t-test. Example blots of cells over-expressing **e**) Flag-HES5 and **f**) Flag-Venus::HES5 treated with 10µM cycloheximide over time (mins). Error bars –SD. Source data are provided in a Source Data file.

Supplementary Figure 2. Quantification of Venus::HES5 in mouse E10.5 spinal cord

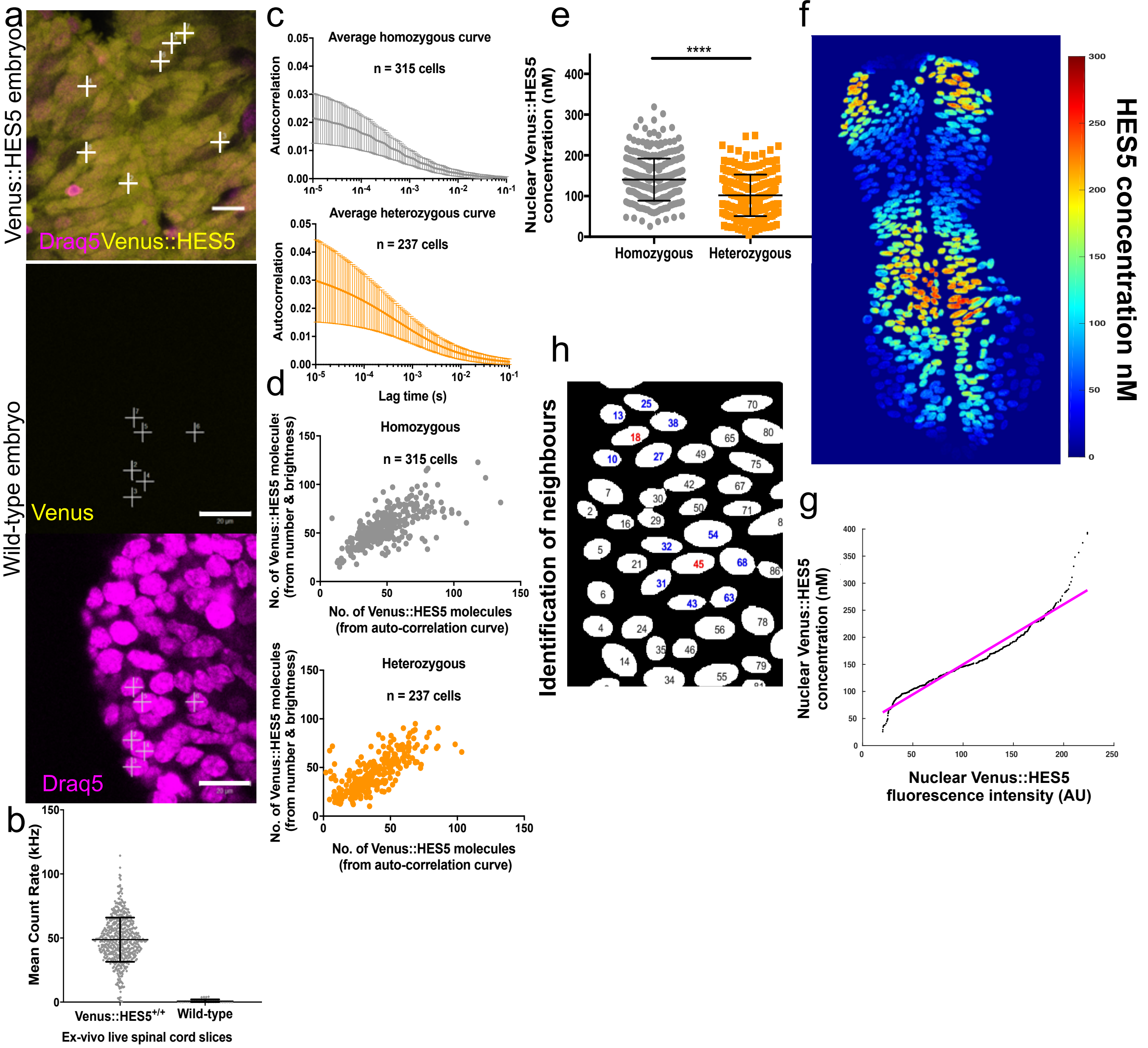


**Supplementary Figure 2. Quantification of Venus::HES5 in mouse E10.5 spinal cord.**
**a**)Snapshot of transverse section of live E10.5 Venus::HES5 (left panel, scale bar 15μm) and wild-type mouse (right panels, scale bar 20μm) ex vivo with live Draq5 nuclear staining. Plus signs mark points of Fluorescence Correlation Spectroscopy recording. **b**) Mean count rates per nucleus from Venus::HES5 homozygous (n=589 cells) and wild-type (n=28 cells) E10.5 spinal cord ex-vivo slices. **c**) Average autocorrelation curves from FCS measurement of 315 cells (4 experiments) in homozygous and 237 cells (4 experiments) in heterozygous Venus::HES5 spinal cord ventral region. **d**) Number of Venus::HES5 molecules in confocal volume calculated from fitting autocorrelation curves plotted against number of molecules from number and brightness analysis of the same single FCS measurement. (Homozygous n=315 cells, r=0.69 and p<0.0001 and Heterozygous n=237 cells, r=0.75 and p>0.0001 in Spearman's rank test). **e**) Nuclear concentration of Venus::HES5 in live E10.5 homozygous (n = 315 cells, 4 experiments) and heterozygous (n = 237, 4 experiments) ex vivo spinal cord slices ventral region (p <0.0001 in two-tailed Mann-Whitney test). **f**) Second example of quantitative map of nuclear Venus::HES5 concentration in live E10.5 ex-vivo spinal cord. Colour bar shows Venus::HES5 concentration by scaling intensity values according to linear fit of Q-Q plot in m**. g**) Q-Q plot of nuclear Venus::HES5 fluorescence intensity vs nuclear Venus::HES5 concentration. **h**) Example image mask showing a cell and its nearest (first rank) neighbours. Manually segmented nuclei in white with nuclei ID number (black). Cells of interest marked in red and nearest neighbours in blue. Error bars –SD. Source data are provided in a Source Data file.

**Supplementary Figure 3. Venus::HES5 fluctuations do not correlate with global changes**

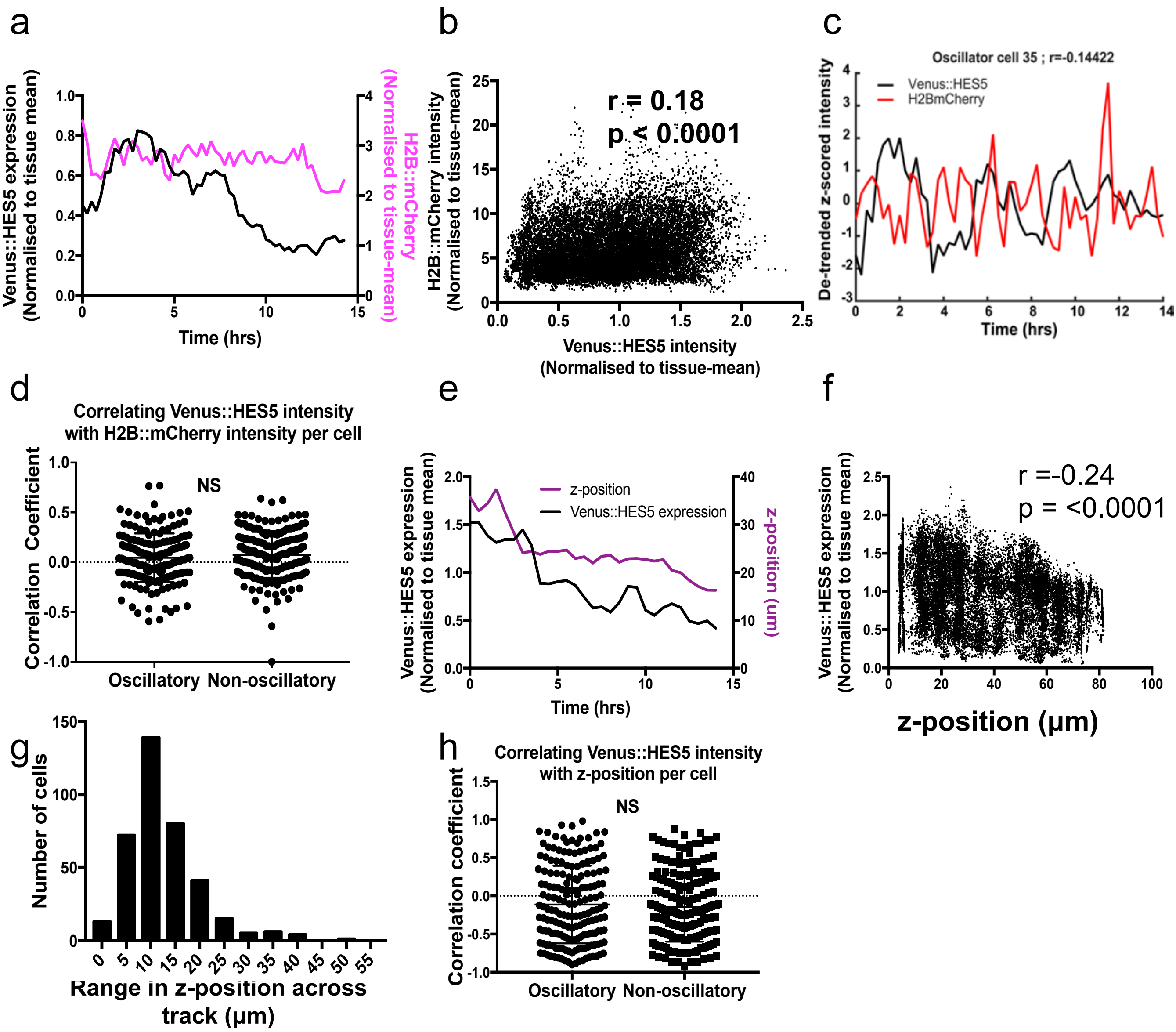


**Supplementary Figure 3. Venus::HES5 fluctuations do not correlate with global changes.**
**a**) Example timeseries of normalised Venus::HES5 intensity and H2B::mCherry intensity in a single cell from an ex-vivo live E10.5 spinal cord slice culture. Single cell intensity values were normalised to the tissue mean intensity over time. **b**) Plot of normalised Venus::HES5 intensity vs normalised H2B::mCherry intensity. All data points from 12-hour tracks of 181 cells in 3 experiments. r = 0.18 from Spearman rank correlation shows no correlation with p <0.0001. **c**) Example timeseries of normalised to tissue mean and de-trended Venus::HES5 and H2B::mCherry intensities in a single cell from an ex vivo live E10.5 spinal cord slice culture. Spearman rank correlation r = -0.14. **d**) Spearman rank correlation coefficient between de-trended Venus::HES5 and de-trended H2BmCherry intensity in single cells classified as having oscillatory (n=180 cells, 3 experiments) or non-oscillatory (n=196 cells, 3 experiments) Venus::HES5 protein expression. Not significant (NS) in Mann-Whitney two-tailed test. **e**) Example timeseries of relative Venus::HES5 intensity and z-position in a single cell from an ex vivo live E10.5 spinal cord slice culture. Single-cell intensity values were normalised to the tissue mean intensity over time. **f**) Plot of z-position vs normalised Venus::HES5 intensity. All data points from 12 hour tracks of 181 cells in 3 experiments. r =-0.24 from Spearman rank correlation shows low correlation with p <0.0001. **g**) Histogram of range in z-positions across single cell tracks over 12 hrs. 181 cells from 3 experiments. Mean=12.4μm, SD=7.5μm. **h**) Spearman rank correlation coefficient between de-trended Venus::HES5 intensity and z-position in single cells classified as having oscillatory (n=180 cells, 3 experiments) or non-oscillatory (n=196 cells, 3 experiments) Venus::HES5 protein expression. Not significant (NS) in Mann-Whitney two-tailed test. Error bars –SD. Source data are provided in a Source Data file.

**Supplementary Figure 4. Hierarchical clustering of Venus::HES5 intensities over time**

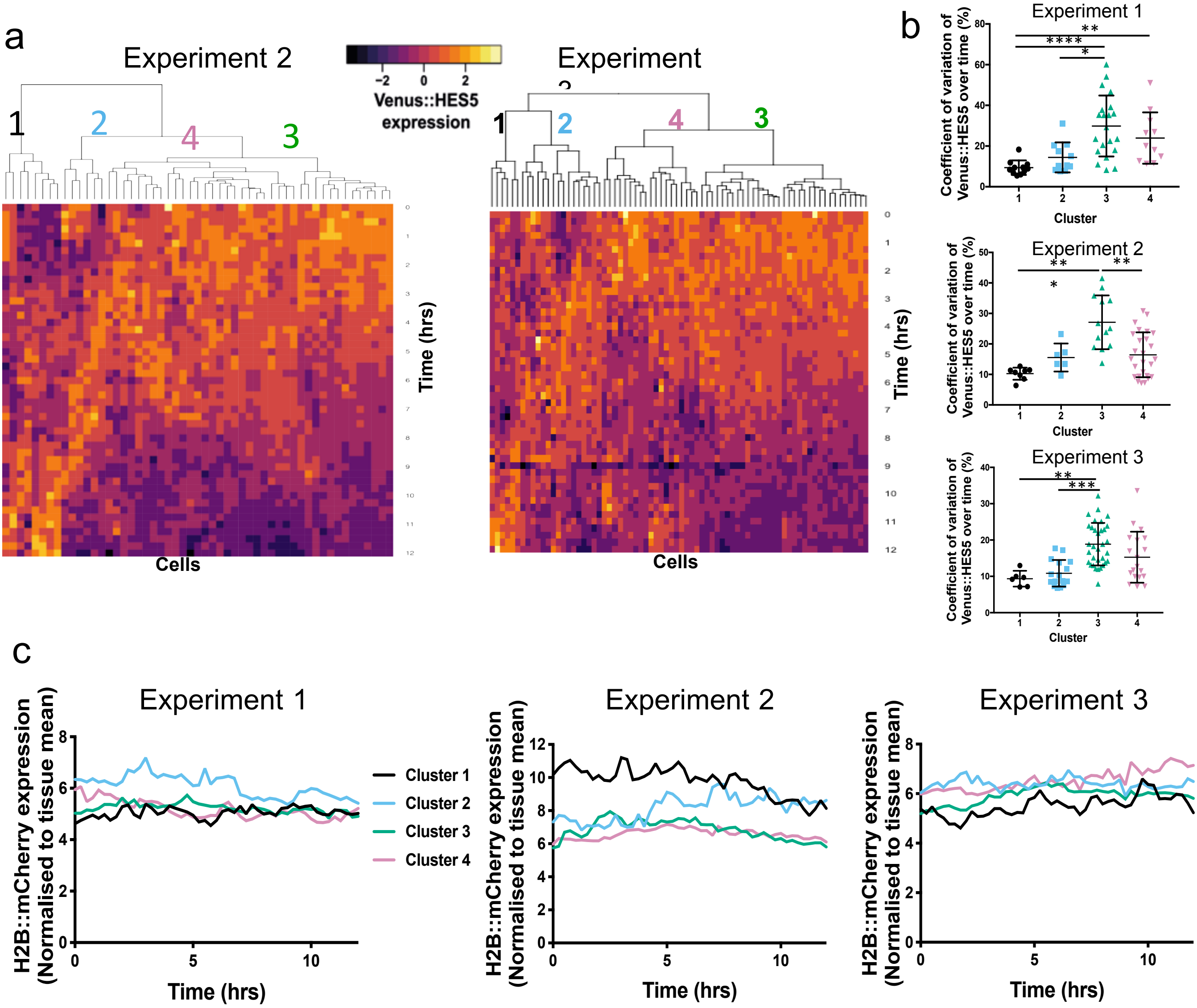


**Supplementary Figure 4. Hierarchical clustering of Venus::HES5 intensities over time.**
**a**) Dendrograms produced by hierarchical clustering of normalised single-cell Venus::HES5 expression dynamics in E10.5 Venus::HES5 Sox1Cre:ERT2 Rosa26RH2BmCherry spinal cord slice culture from two independent experiments (Experiment 2 and 3) conducted in the same conditions as data shown in **Fig. 2h** (Experiment 1). Columns show individual cell Venus::HES5 expression dynamics in a heatmap aligned to start at t=0, the start of tracking. Rows represent time points. All cells tracked for 12-hour time window with 15 minute frame intervals. **b**) Coefficient of variation of Venus::HES5 expression in a single cell over time was used to identify and annotate corresponding clusters in Experiments 1 to 3. Error bars -SD. Kruskal-Wallis with Dunn's multiple comparison test used to define significance levels. '*' indicates p<0.05, '**' indicates p<0.01, '***' indicates p<0.001, '***' indicates p<0.0001. **c**) Mean H2B::mCherry expression dynamics for cells in each cluster in each experiment. (Cell number per experiment, Cluster 1 – 11, 8, 6 cells, cluster 2 – 11, 6, 16 cells, cluster 3 – 21, 13, 33 cells, cluster 4 – 11, 26, 19 cells. Source data are provided in a Source Data file.

**Supplementary Figure 5. Inferring cell fate from positional information**

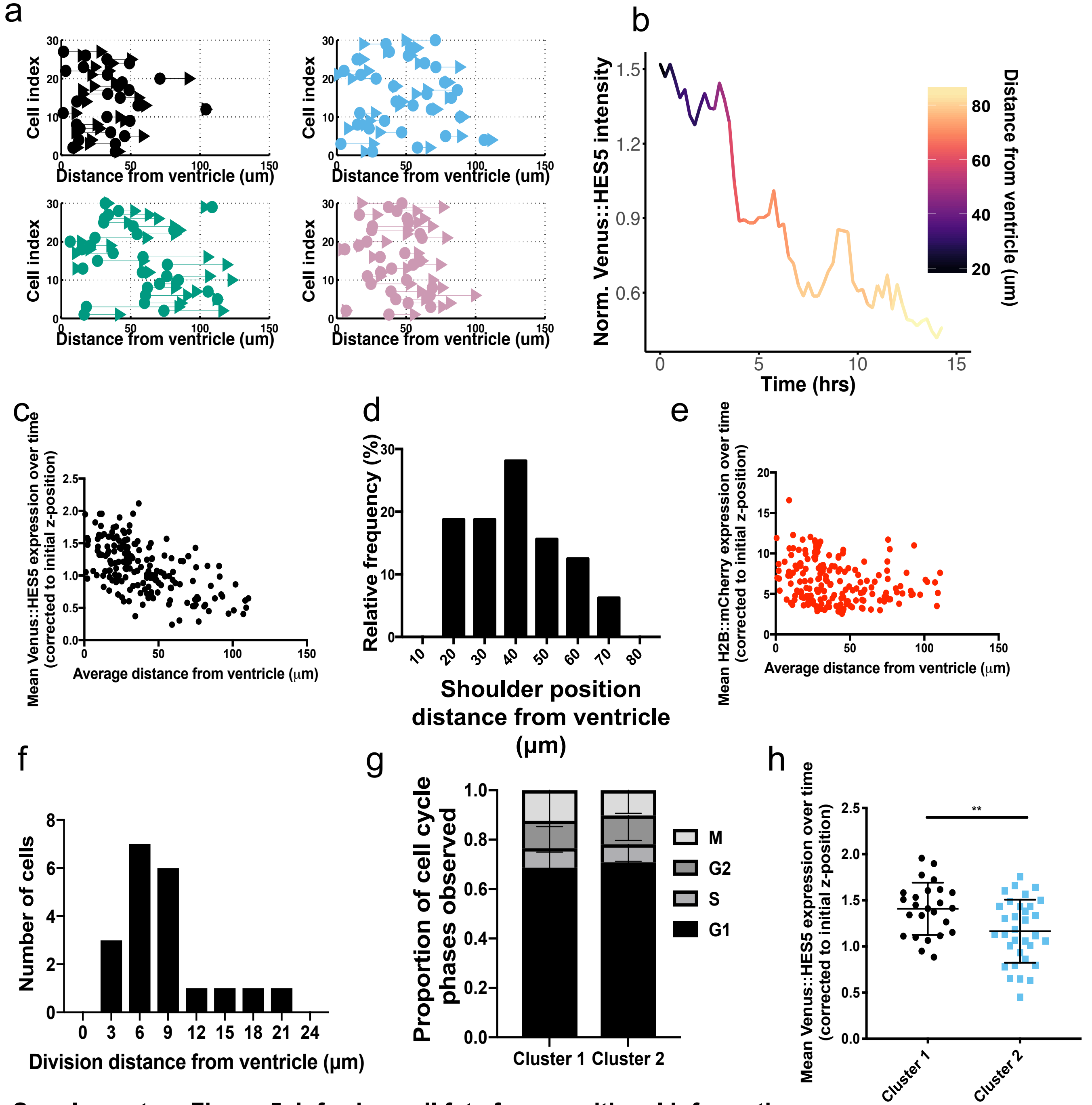


**Supplementary Figure 5. Inferring cell fate from positional information.**

**a**) Displacement of single cells in each cluster in apico-basal axis, measured from ventricle as 0. Dot represents start position and arrow represents finish position of each cell. 181 cells from 3 experiments clustered separately. **b)** Single cell Venus::HES5 normalised intensity from cluster 3 colour-coded by distance from ventricle in apico-basal axis. **c**) Average distance from the ventricle over 12 hr track vs mean Venus::HES5 corrected intensity. Intensity was corrected for the increased light scattering at increasing depth (z-position) in to tissue by normalising to initial z-position in tissue (see methods). 181 cells from 3 experiments. **d**) Distribution of distance from the ventricle at which cells in cluster 3 and 4 start to turn off Venus::HES5 (shoulder point). mean=40.3μm, SD=14.6μm. n=32 cells, 3 experiments. **e**) Average distance from the ventricle over 12 hr track vs mean H2B::mCherry intensity corrected for z-position in the tissue. 181 cells from 3 experiments. **f**) Distribution of distance from the ventricle at which cells in cluster 1 & 2 divide. mean=8.8μm, SD=4.9μm. n= 20 cells, 3 experiments. **g**) Proportion of cell-cycle phases observed in cells of cluster 1 vs cluster 2. **h**) Mean Venus::HES5 intensity corrected for z-position in the tissue in cells of cluster 1 vs cluster 2. Student t-test p=0.005 (**). Error bars - SD. Source data are provided in a Source Data file.

Supplementary Figure 6. Inferring cell fate from Notch inhibition.

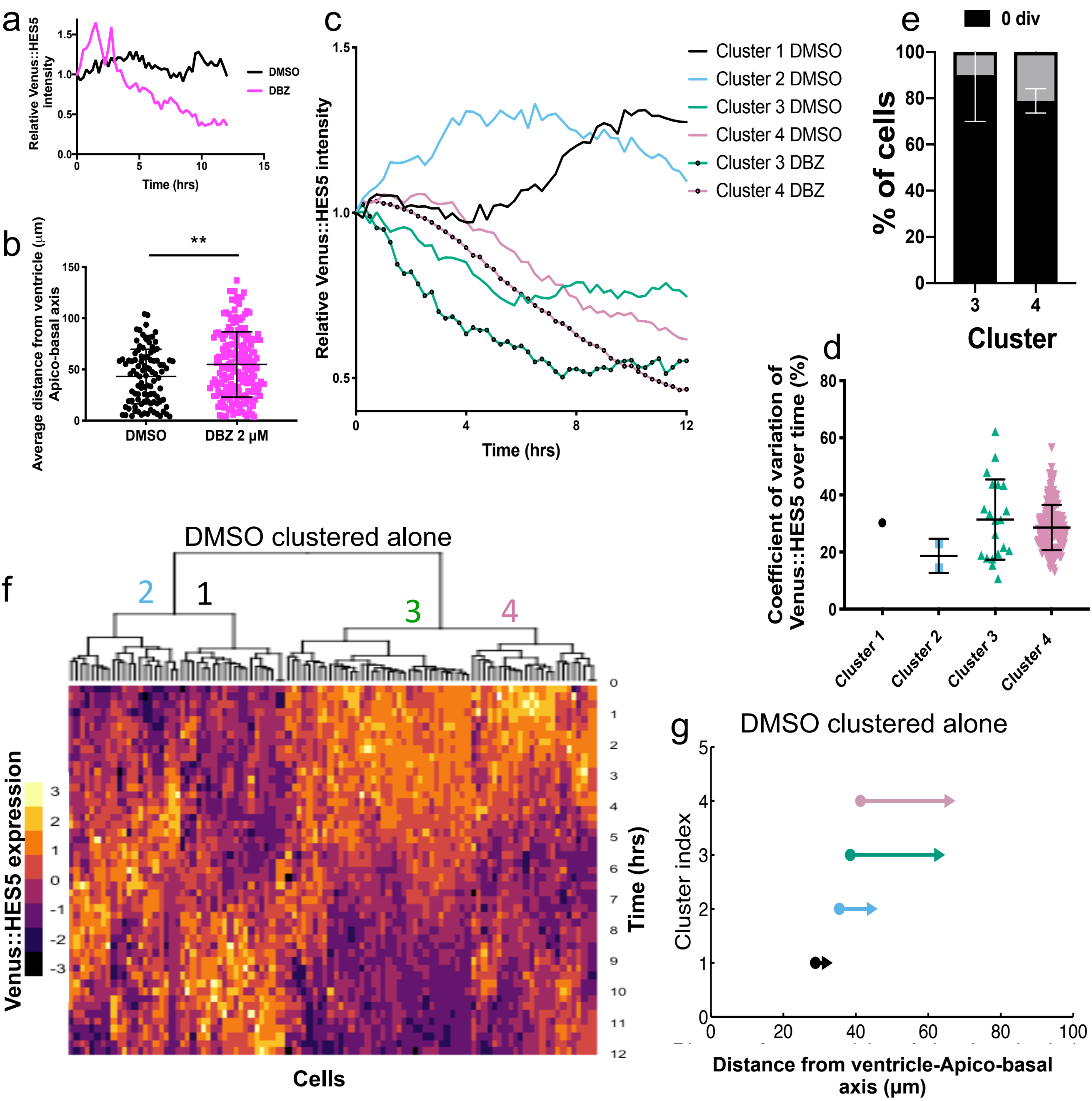


**Supplementary Figure 6. Inferring cell fate from Notch inhibition.**
**a**) Example relative Venus::HES5 expression in DMSO and 2μM DBZ treated ex vivo slices. **b**) Average distance from ventricle (μm) in single cell observed over 12-hour track measured in apico-basal axis in DMSO and 2μM DBZ treated E10.5 ex vivo spinal cord slices. n=4 experiments. p=0.0036(**) in two-tailed Mann-Whitney test. **c**) Mean Venus::HES5 expression dynamics for cells in each cluster in DMSO and 2μM DBZ treated ex vivo slices. Cells in 4 experiments clustered together (Cell number, DMSO Cluster 1 – 17 cells, cluster 2 – 24 cells, cluster 3 – 27 cells, cluster 4 – 33 cells. DBZ Cluster 3 – 20 cells, cluster 4 – 171 cells. **d**) Coefficient of variation of Venus::HES5 expression in a single cell over time. **e**) Percentage of cells showing a division in 12 hour track in 2μM DBZ treated ex vivo slices. n=3 experiments. **f**) Hierarchical clustering of standardized single-cell Venus::HES5 expression in DMSO treated E10.5 ex-vivo slices. 100 cells in 3 experiments clustered together. Columns show single cell standardised Venus::HES5 expression dynamics in a heatmap. All cells tracked for 12-hour time window with 15-minute frame intervals. **g**) Displacement of cells in DMSO treated E10.5 ex vivo spinal cord slides in each cluster. Dot represents average start position and arrow represents average finish position of cells in each cluster. DMSO n=100 cells, 2μM DBZ n=195 cells, 3 experiments. Error bars – SD. Source data are provided in a Source Data file.

**Supplementary Figure 7. Single cell Venus::HES5 expression in cells from clusters 1 to 4**

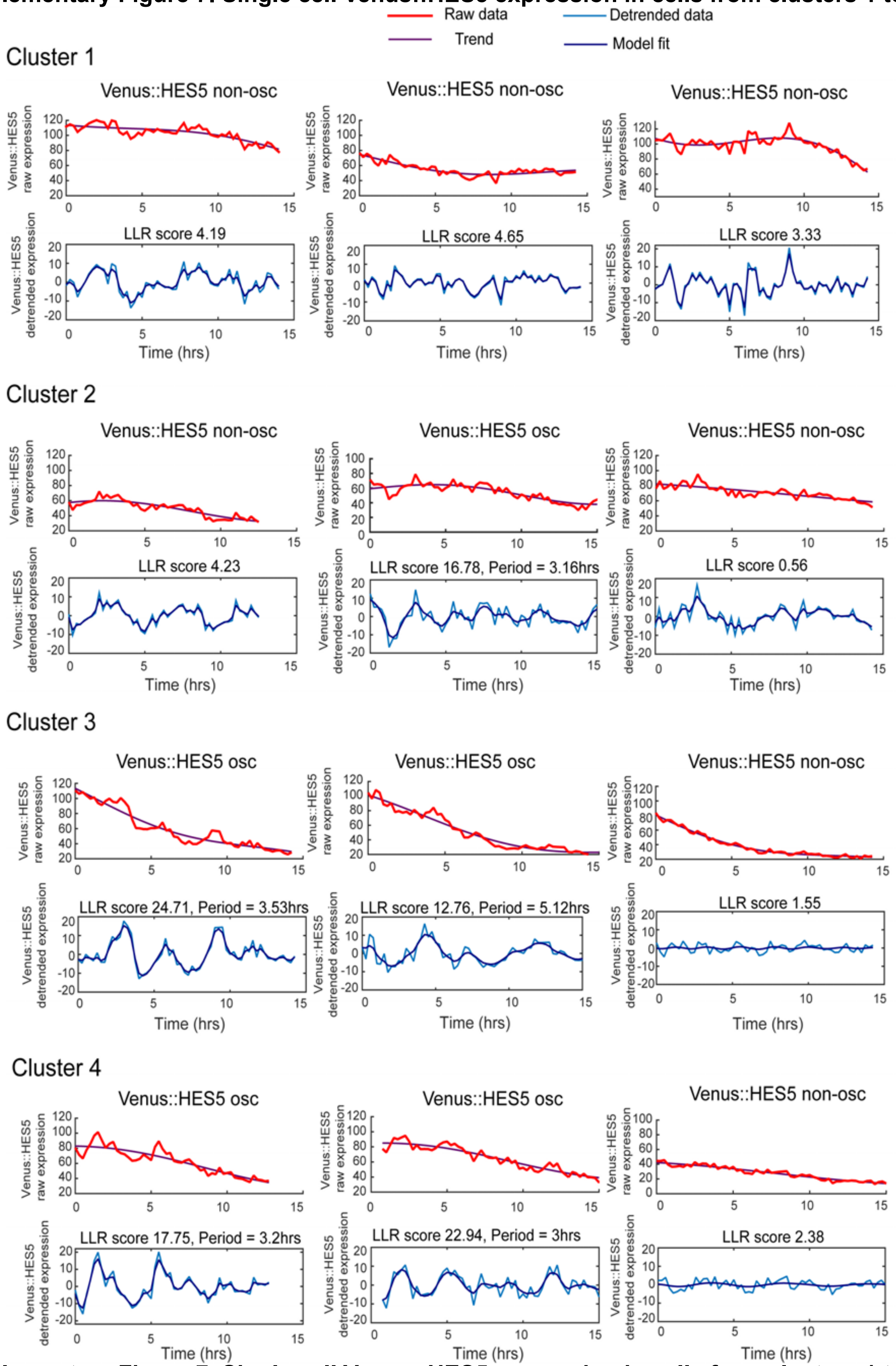


**Supplementary Figure 7. Single cell Venus::HES5 expression in cells from clusters 1 to 4.** Single cell timeseries of Venus::HES5 expression in ex vivo live E10.5 Venus::HES5$^{+/-}$ spinal cord slice cultures in cells from cluster 1-4. Labelled as having oscillatory (osc.) or non-oscillatory expression Top panel - raw Venus::HES5 intensity (before normalisation to tissue mean intensity) in red, with trend in purple. Trend subtracted to give de-trended data in lower panel (blue) with covariance model fit (purple). Source data are provided in a Source Data file.

**Supplementary Figure 8. Statistical modelling and detection of oscillatory Venus::HES5**

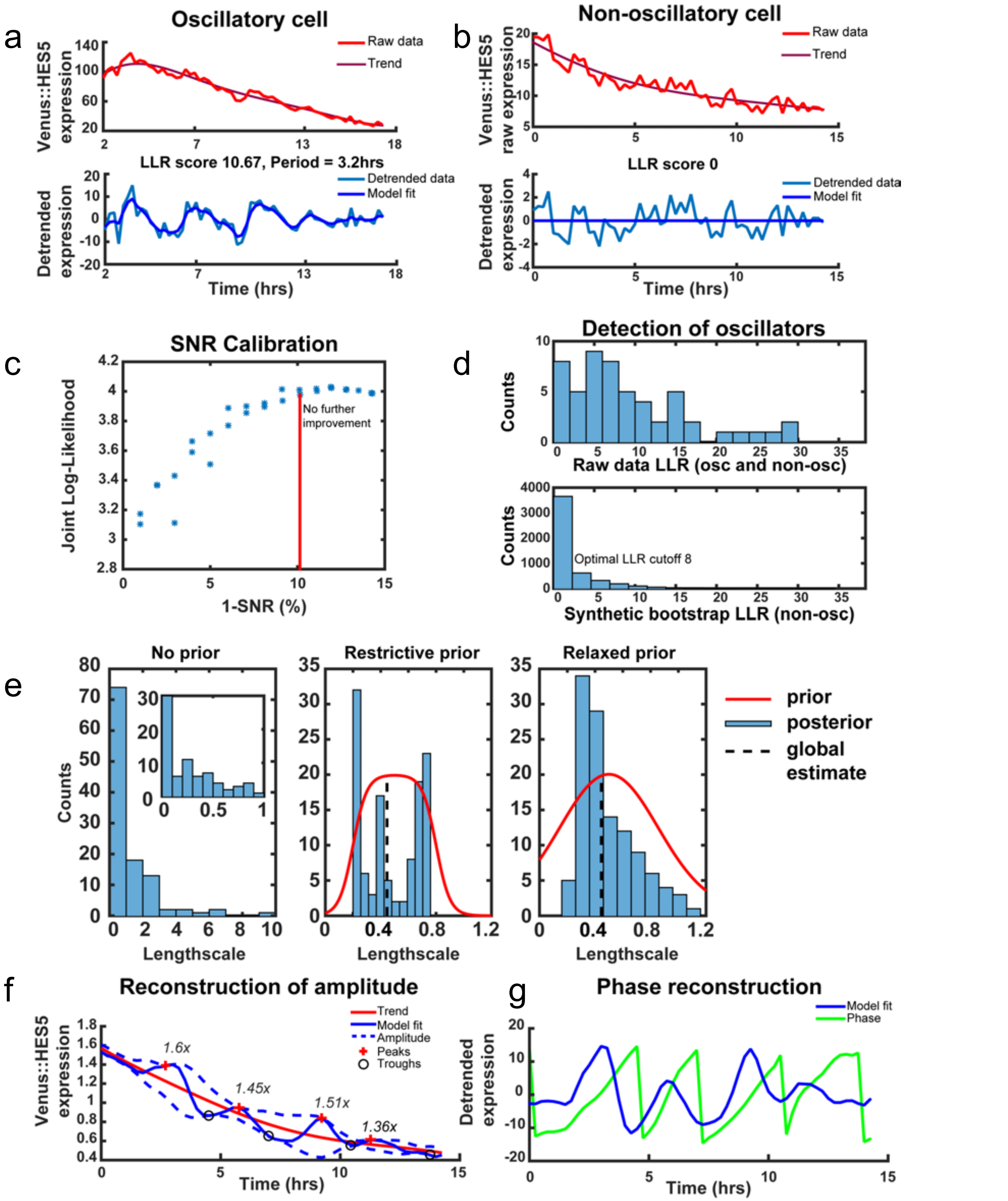


**Supplementary Figure 8. Statistical modelling and detection of Venus::HES5.**
Example of **a**) oscillatory and **b**) non-oscillatory Venus::HES5 in spinal cord ex-vivo slices; intensity traces corrected by tissue mean and fitting of long term trend (top panel); detrended data and short term trend fitted with the OUosc covariance model (bottom panel); **c**) Evaluated joint log-likelihood as a function of signal-to-noise ratio (SNR) shows saturation of log-likelihood values for noise ≥10.14%; **d**) distribution of LLR scores in spinal cord tissue real data vs synthetic non-oscillatory data and optimal LLR threshold at 3% FDR; **e**) estimation of lengthscale: without a prior (panel 1) where insert shows the distribution of values between 0 and 1; using the smooth box1 prior with parameters *l=0.2;L=0.8; η=20* (panel 2) and *l=0.2;L=0.8; η=5* (panel 3). **f**) Hilbert reconstruction of amplitude of Venus::HES5 expression **g**) phase reconstruction of detrended Venus::HES5. Peaks (red plus sign) and troughs (black circle) identified from phase.

Supplementary Figure 9. Venus::HES5 expression dynamics in primary neural progenitors

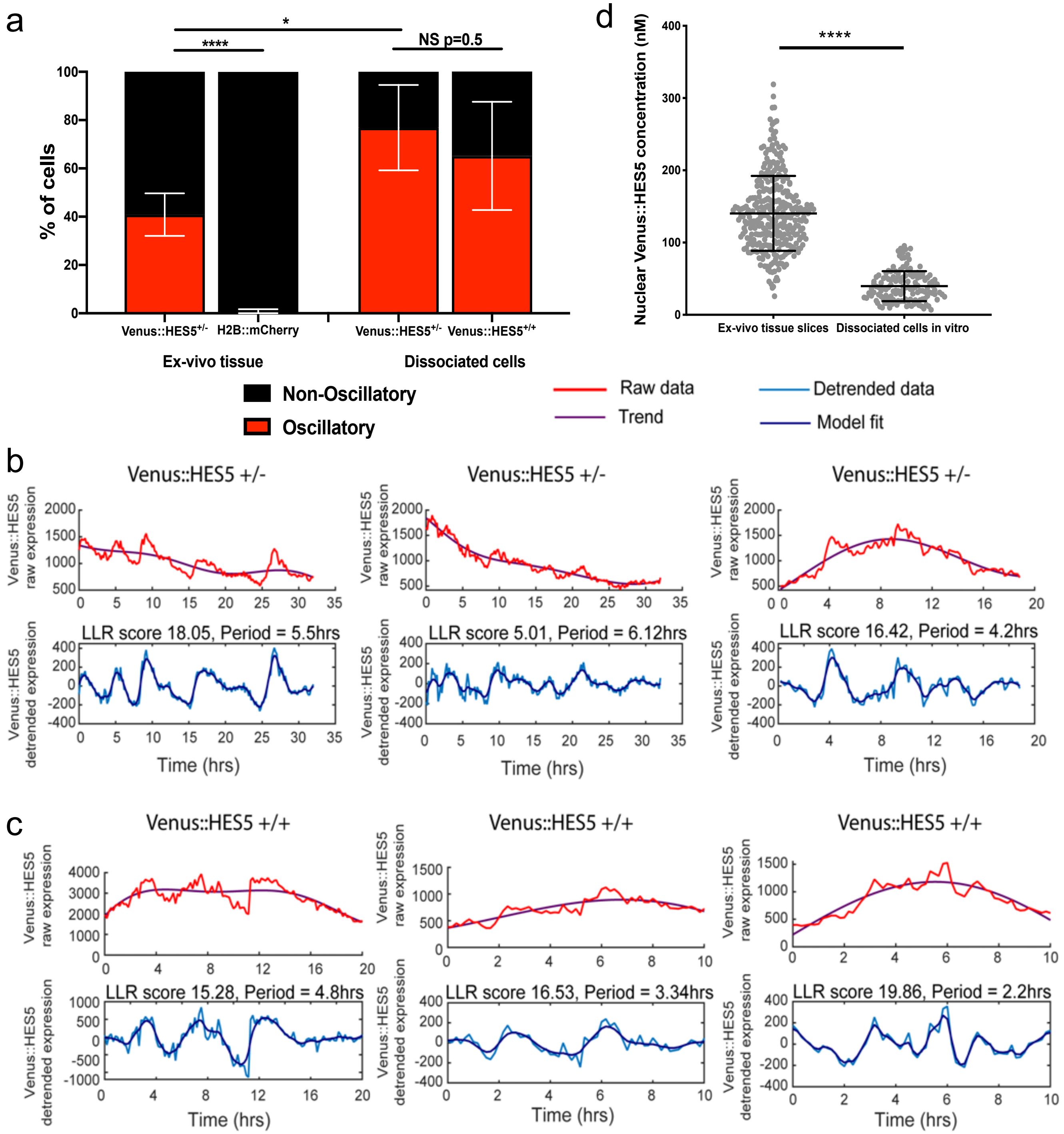


**Supplementary Figure 9. Venus::HES5 expression dynamics in primary neural progenitors.**

**a)** Percentage of cells with oscillatory Venus::HES5 or H2B::mCherry expression dynamics in heterozygous ex-vivo spinal cord slices and heterozygous vs homozygous dissociated cells cultured in vitro. Ex-vivo data 181 cells, 3 experiments. Frequency data analysed in two-sided Fisher's exact test $p<0.0001$. Dissociated cells Venus::HES5$^{+/-}$ (n=60 cells) vs Venus::HES5$^{+/+}$ (n=35 cells) 4 experiments. NS – no significance in Student t-test. Ex-vivo Venus::HES5$^{+/-}$ vs dissociated cells Venus::HES5$^{+/-}$ p=0.02 in Student t-test. **b-c)** Single cell Venus::HES5 expression in dissociated primary neural progenitor cells from spinal cord of E10.5-E11.5 **b**) Venus::HES5$^{+/-}$ embryos and **c**) Venus::HES5$^{+/+}$ embryos. Top panel of each shows Venus::HES5 intensity in red, with trend in purple. The trend was subtracted to give de-trended data in lower panel (blue) shown with covariance model fit (purple). **d**) Nuclear Venus::HES5$^{+/+}$ concentration from FCS in ex-vivo tissue slices from E10.5 Venus::HES5$^{+/+}$ embryos (n=315 cells) and dissociated primary neural stem cells (n=139 cells). Two-tailed Mann-Whitney test $p<0.0001$(****). Error bars show SD. Source data are provided in a Source Data file.

**Supplementary Figure 10. Approximate Bayesian Computation to parameterise *Hes5* model**

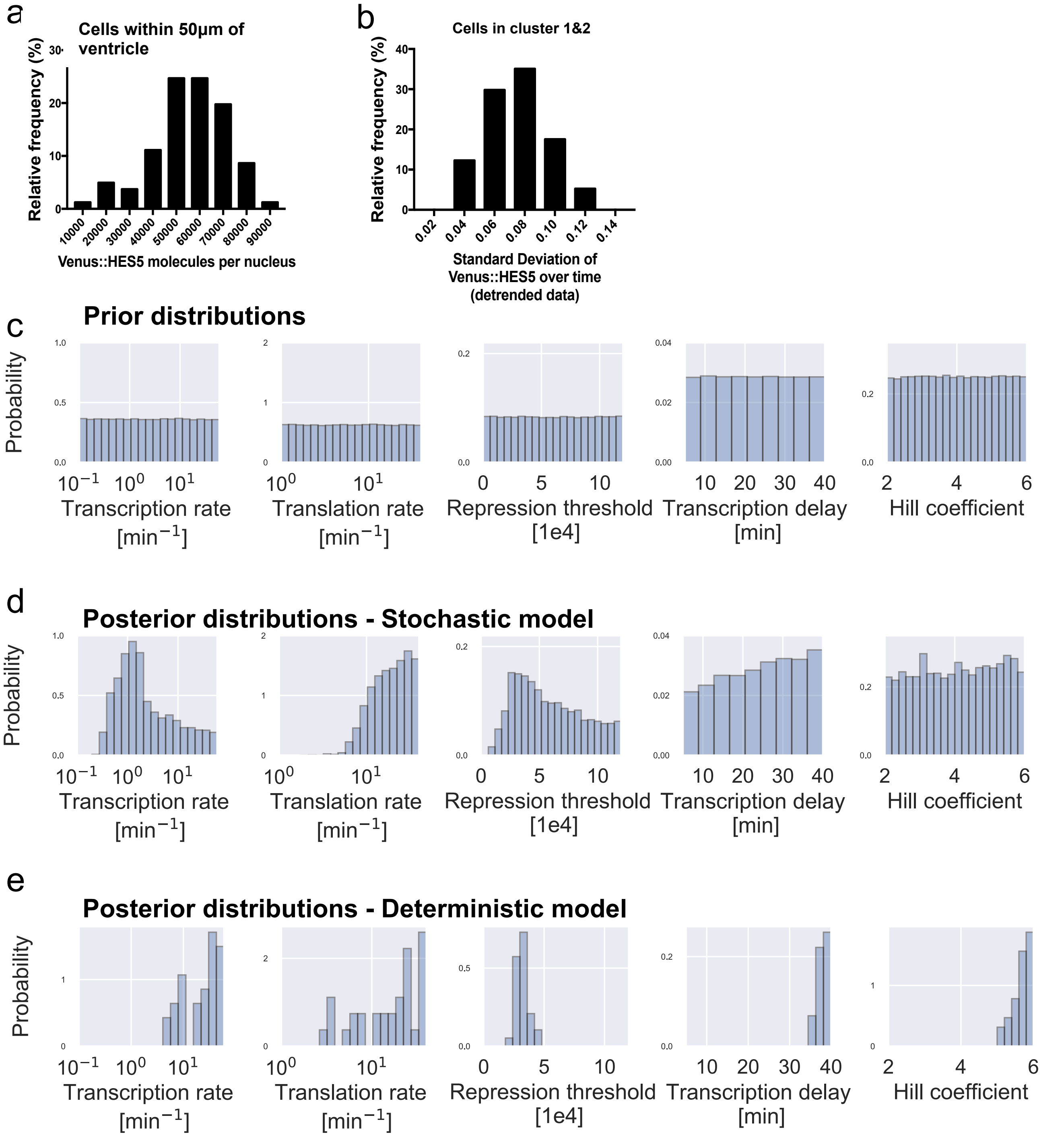


**Supplementary Figure 10. Approximate Bayesian Computation approach to parameterise *Hes5* model. a**) Distribution of Venus::HES5 molecule number per nucleus from cells less than 50um from ventricle in quantitative map of Venus::HES5$^{+/+}$ expression shown in Fig.1j. Mean=56,600 SD=15,000 n= 81 cells. **b**) Distribution of the single-cell standard deviation of Venus::HES5 expression over time from detrended data. 57 cells in clusters 1 and 2 from 3 experiments. Mean=0.074, SD=0.02. **c**) Prior distributions (200000 samples) and **d**) posterior distributions of individual model parameters of the stochastic *Hes5* auto-negative feedback model (4901 samples). Posterior distributions can be interpreted as the probability of the parameters given the data. Inferred model parameters are associated with high uncertainty. **e**) Posterior distributions of individual model parameters of the deterministic auto-negative feedback model. 32 out of 200,000 parameter points tested in the deterministic model are accepted with criteria of 55,000 to 65,000 molecule number and >5% standard deviation in HES5 levels over time.

**Supplementary Figure 11. Reduction in Venus::HES5 levels does not cause oscillations**

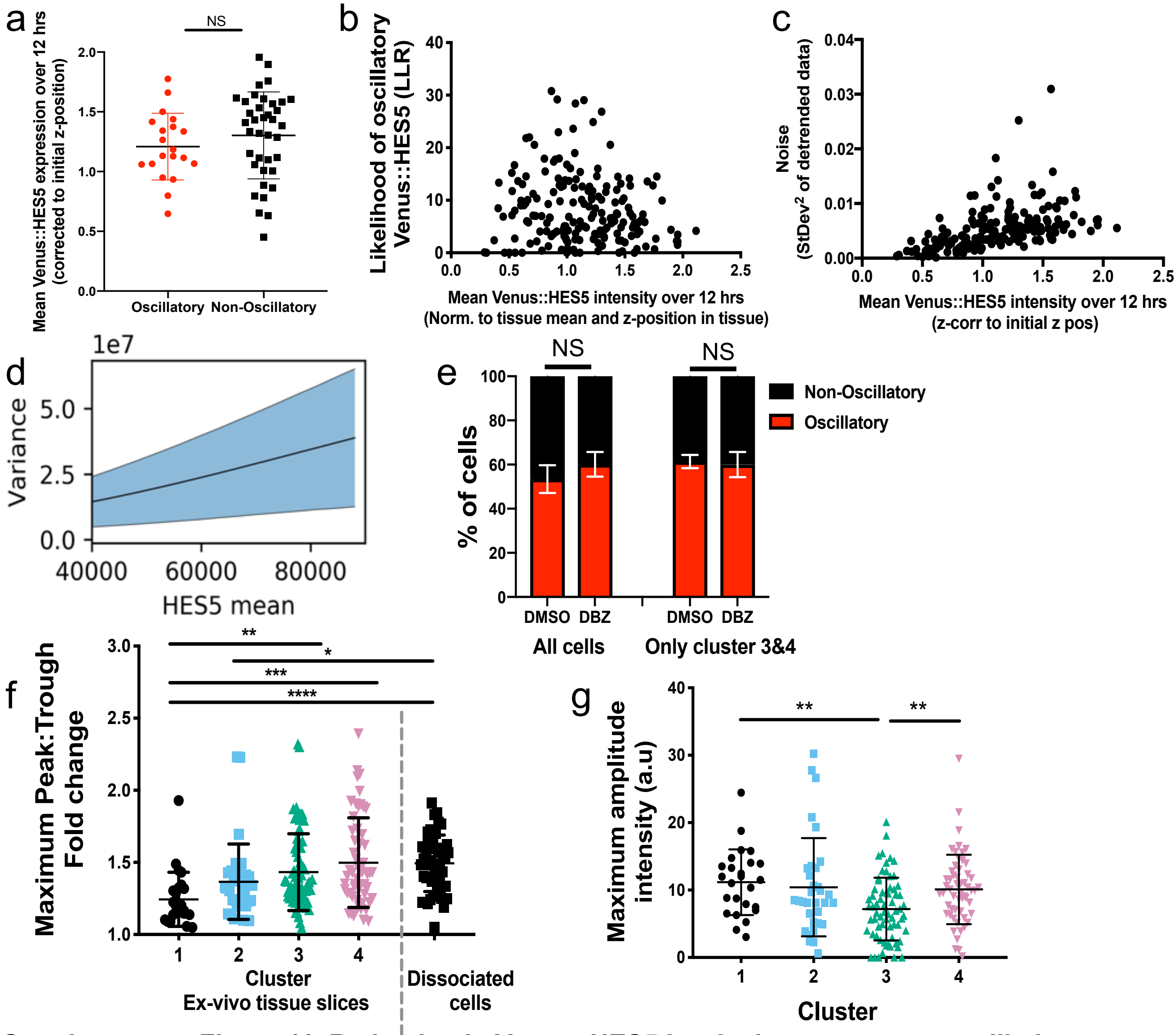


**Supplementary Figure 11. Reduction in Venus::HES5 levels does not cause oscillations.**
**a**) Corrected mean Venus::HES5 intensity of oscillatory (n=20) or non-oscillatory (n=38) cluster 1 & cluster 2 cells. 3 experiments clustered separately. NS – non-significant in Student t-test. **b**) Corrected mean Venus::HES5 intensity vs likelihood of oscillatory Venus::HES5 (LLR from stochastic test). 181 cells, 3 experiments. **c**) Corrected mean Venus::HES5 intensity vs Venus::HES5 noise (calculated as $SD^2$ of detrended Venus::HES5 intensity). 181 cells, 3 experiments. **d**) Bayesian posterior prediction for the relationship between HES5 mean and absolute variance when changing HES5 repression threshold. Black line shows posterior mean and blue area shows SD. **e**) Percentage of cells with oscillatory Venus::HES5 expression dynamics in heterozygous ex-vivo spinal cord slices treated with DMSO or 2μM DBZ, showing all cells (DMSO n=103 cells, DBZ n=195 cells, 4 experiments ) or just cluster 3&4 cells (DMSO n=61 cells, DBZ n=192 cells, 4 experiments). NS - No significance in Student t-test. **f**) Maximum peak-to-trough fold-change in Venus::$HES5^{+/-}$ expression ex-vivo per cluster and in dissociated primary neural progenitor cells. Kruskal-Wallis with Dunn's multiple comparison test shows cluster 1 vs 3 adjusted p=0.0037 (**), cluster 1 vs 4 adjusted p=0.0002 (***), 1 vs dissoc. cells adjusted p<0.0001 (****), cluster 2 vs dissoc. cells adjusted p=0.02 (*). 181 cells, 3 experiments clustered separately. Cluster 1 n=27 cells, cluster 2 n=33 cells, cluster 3 n=67 cells, cluster 4 n=54 cells, dissociated cells n=50. **g**) Maximum amplitude in single cell Venus::$HES5^{+/-}$ expression in ex-vivo slices per cluster. Amplitude calculated after the trend in Venus::HES5 expression is subtracted. Kruskall-Wallis with Dunn's multiple comparisons test indicated cluster 1 vs 3 adjusted p-value = 0.0043 (**), cluster 3 vs 4 adjusted p-value = 0.0067 (**). 181 cells, 3 experiments clustered separately. Error bars – SD. Source data provided in Source Data file.

**Supplementary Figure 12. Venus::HES5 expression in DMSO and DBZ treated ex-vivo slices**

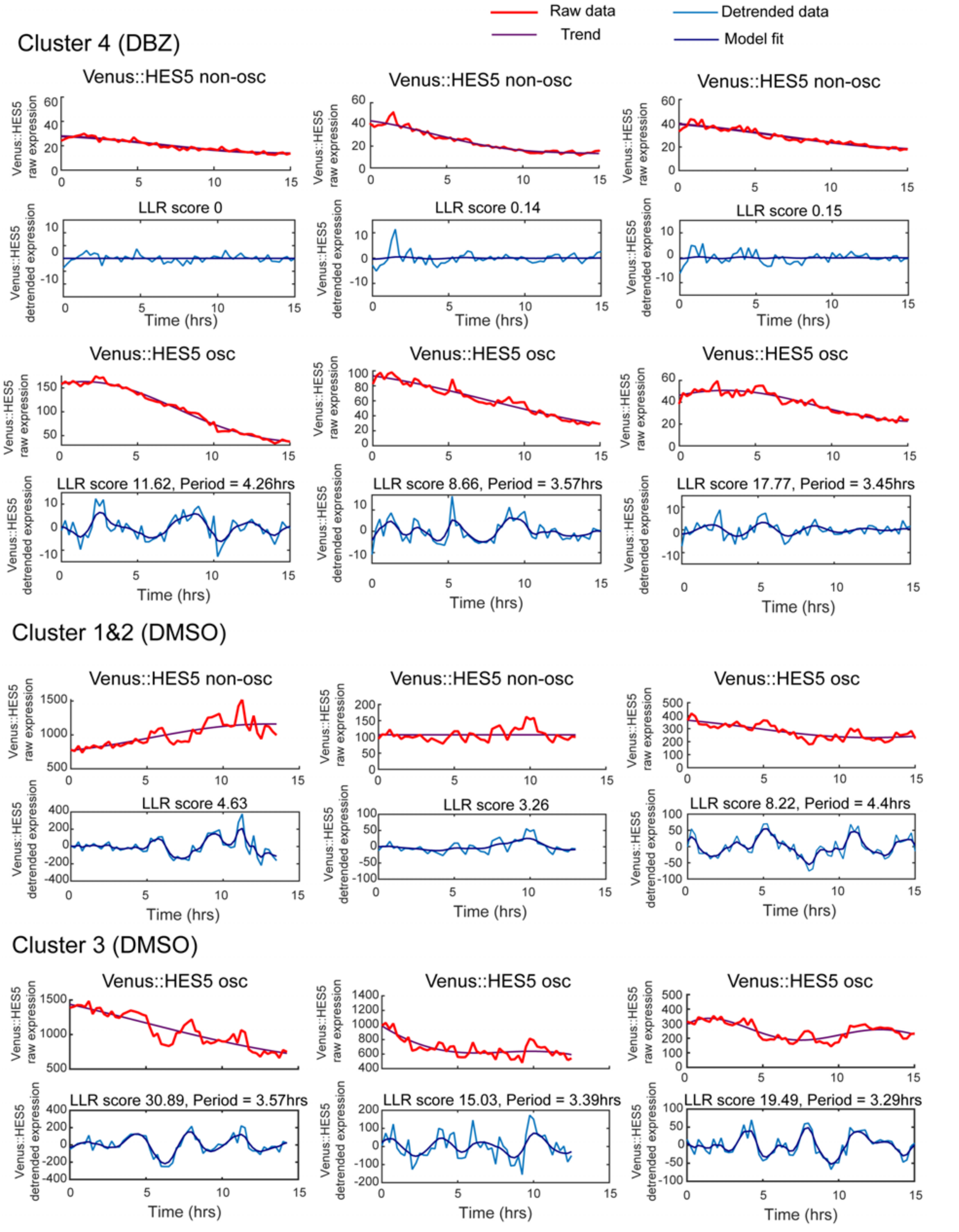


**Supplementary Figure 12. Venus::HES5 expression in DMSO and DBZ treated ex-vivo slices.** Single cell timeseries of Venus::HES5 protein expression in ex vivo live E10.5 Venus::HES5$^{+/-}$ spinal cord slice cultures treated with DMSO or 2µM DBZ. Example cells labelled as having oscillatory (osc.) or non-oscillatory (non-osc) Venus::HES5 expression. Top panel of each shows raw Venus::HES5 intensity (before normalisation to tissue mean intensity) in red, with trend in purple. The trend was subtracted to give de-trended data in lower panel (blue) shown with covariance model fit (purple). DMSO or DBZ was added immediately prior to imaging. Source data are provided in a Source Data file.

### Supplementary Table 1 : Model implementation and inference parameters

| Parameter | Description | Value |
|---|---|---|
| $\Delta t$ | Time step | 1 min |
| $t_{\mathrm{eq}}$ | Equilibration time | 1000min |
| $t_{\mathrm{obs}}$ | Observation time | 7500min |
| $P_{\mathrm{in}}$ | Initial protein number | $P_0$ |
| $M_{\mathrm{in}}$ | Initial mRNA number | 10 |
| $N_P$ | Number of simulations per parameter point | 200 |
| $N_{\mathrm{tot}}$ | Number of prior samples | 200000 |

### Supplementary Table 2 : Model parameters

| Parameter | Description | Value | Method | Reference |
|---|---|---|---|---|
| $\alpha_m$ | Basal transcription rate | 0.01/min - 60/min | Bayesian inference | (Singh and Padgett, 2009; Suter et al., 2011) |
| $\alpha_p$ | Translation rate | 1/min - 40/min | Bayesian inference | (Alberts et al., 2008; Boström et al., 1986; Ingolia et al., 2011; Schwanhäusser et al., 2011) |
| $P_0$ | Repression threshold | 0 - 120000 | Bayesian inference | (Monk, 2003), see methods text |
| $\tau$ | Time delay | 5 min - 40 min | Bayesian inference | (Lewis, 2003; Monk, 2003; Phillips et al., 2016) |
| $n$ | Hill coefficient | 2 - 6 | Bayesian inference | (Galla, 2009; Monk, 2003; Phillips et al., 2016) |
| $\mu_m$ | mRNA degradation rate | log(2)/(30min) | Measurement | - |
| $\mu_p$ | Protein degradation rate | log(2)/(90min) | Measurement | - |
